# Spliformer-V2 enables multi-tissue prediction and interpretation of splice-altering genetic variants

**DOI:** 10.1101/2025.06.28.662167

**Authors:** Xuelin Tang, Meiyu Shao, Helei Lei, Xingyu Ma, Jingyan Guo, Yifan Shen, Qihui Wu, Yi Dong, Yi Zeng, Aaron D Gitler, Yelin Chen, Agessandro Abrahão, Lorne Zinman, Ekaterina Rogaeva, Yan Chen, Justin K. Ichida, Ming Zhang

## Abstract

Precise regulation of pre-mRNA splicing underlies transcriptomic diversity and is disrupted in aging and disease, yet tissue-specific splice-altering genetic variants remain poorly resolved. Here, we present Spliformer-V2, a SegmentNT-based deep learning model for predicting and interpreting variant effects on RNA splicing across human tissues. We generated a diploid sequence resolved RNA splice map from paired whole-genome-sequencing and RNA-seq data across 12 central nervous system (CNS) and 6 peripheral tissues for model development. Spliformer-V2 outperformed SpliceTransformer, Pangolin and AlphaGenome in predicting splice-site usage, identified tissue-specific splicing regulatory motifs, and revealed tissue vulnerability to pathogenic splice-altering variants. Analyses of loci associated with 8 neurological diseases prioritized CNS-specific mis-splice-vulnerable genes. In 1,405 amyotrophic lateral sclerosis (ALS) genomes, Spliformer-V2 nominated rare splice-altering variants enriched in *PTPRN2*, which showed reduced expression in TDP-43-depleted neurons. *PTPRN2* overexpression rescued *C9ORF72*-patient derived motor neuron degeneration and modulated TDP-43 mislocalization, indicating it as a potential therapeutic modifier in ALS.

## Introduction

Precise regulation of pre-mRNA splicing generates transcriptomic and proteomic diversity essential for cellular identity, tissue function and organismal homeostasis ^1^. Splicing disruption contributes broadly to human disease, including neurodevelopmental and neurodegenerative disorders ^2^. Although canonical splice-site mutations explain a subset of Mendelian diseases, most disease-associated variants identified by genome-wide association studies (GWAS) reside in noncoding regions, where their functional consequences remain incompletely understood and may act through RNA processing, including alternative splicing ^3, 4^.

A major challenge in interpreting noncoding variation is the tissue specificity of splicing regulation, as the same variant may have different effects across tissues owing to differences in RNA-binding proteins or chromatin context ^5, 6^. This is particularly important in neurological diseases with region-selective pathology. For example, amyotrophic lateral sclerosis (ALS) preferentially affects motor cortex and spinal cord, whereas Alzheimer’s disease (AD) and Parkinson’s disease (PD) show distinct vulnerability across cortical and subcortical regions. In ALS and related TDP-43 proteinopathies, nuclear depletion of TDP-43 drives cryptic exon inclusion, reducing expression of neuronal genes such as *UNC13A* and *STMN2* ^7–10^, suggesting that RNA mis-splicing may contribute to selective neuronal vulnerability. However, most splice-prediction frameworks lack CNS subregion resolution, leaving tissue-specific effects of rare variants and noncoding GWAS loci unresolved.

Deep learning models such as SpliceAI ^11^ and Spliformer ^12^ predict constitutive splice-site disruption with high accuracy, whereas tissue-aware models, including Pangolin ^13^ and SpliceTransformer ^14^, incorporate multi-tissue context. However, existing models remain limited by sparse tissue-specific training data, incomplete representation of CNS subregions or insufficient modeling of genotype-dependent effects. These limitations restrict their utility for interpreting disease-associated noncoding variants in anatomically selective neurological disorders.

Recent DNA foundation models provide an opportunity to address these gaps. Trained on large genomic corpora, these models (such as Nucleotide Transformer) learn transferable representations of regulatory sequence grammar ^15, 16^. However, applying them to splicing biology requires paired genome–transcriptome datasets that link individual genetic variation to tissue-resolved splice-site usage, particularly across CNS regions that are difficult to obtain but central to neurological disease.

Here we present Spliformer-V2, a deep learning framework for predicting variant effects on RNA splicing across 18 human tissues. Spliformer-V2 takes diploid sequences as input, enabling modeling of heterozygous and homozygous variant effects, and predicts single-nucleotide-resolution splice-site usage across tissues. Spliformer-V2 outperforms existing models, improves sQTLs and rare splice-altering variants interpretation, reveals tissue-specific splicing regulatory motifs, and prioritizes mis-splice vulnerable disease genes. Analyses of ALS genomics nominate *PTPRN2* as a splice-altering hot spot gene and a therapeutic modifier that rescues disease motor neuron degeneration and modulates TDP-43 mislocalization.

## Results

### Spliformer-V2 architecture and a diploid sequence-resolved multi-tissue RNA splice map

We developed Spliformer-V2, a multi-tissue deep learning model for single-nucleotide-resolution prediction of splice-site usage (**Fig. 1A**). Spliformer-V2 fine-tunes SegmentNT^16^, which incorporates the DNA encoder from Nucleotide Transformer (v2-500M multi-species) ^15^, pretrained on more than 10.24 billion nucleotide tokens, and replaces the masked-language-modeling head with a 1D U-Net segmentation module nucleotide-level prediction.

**Fig. 1.**
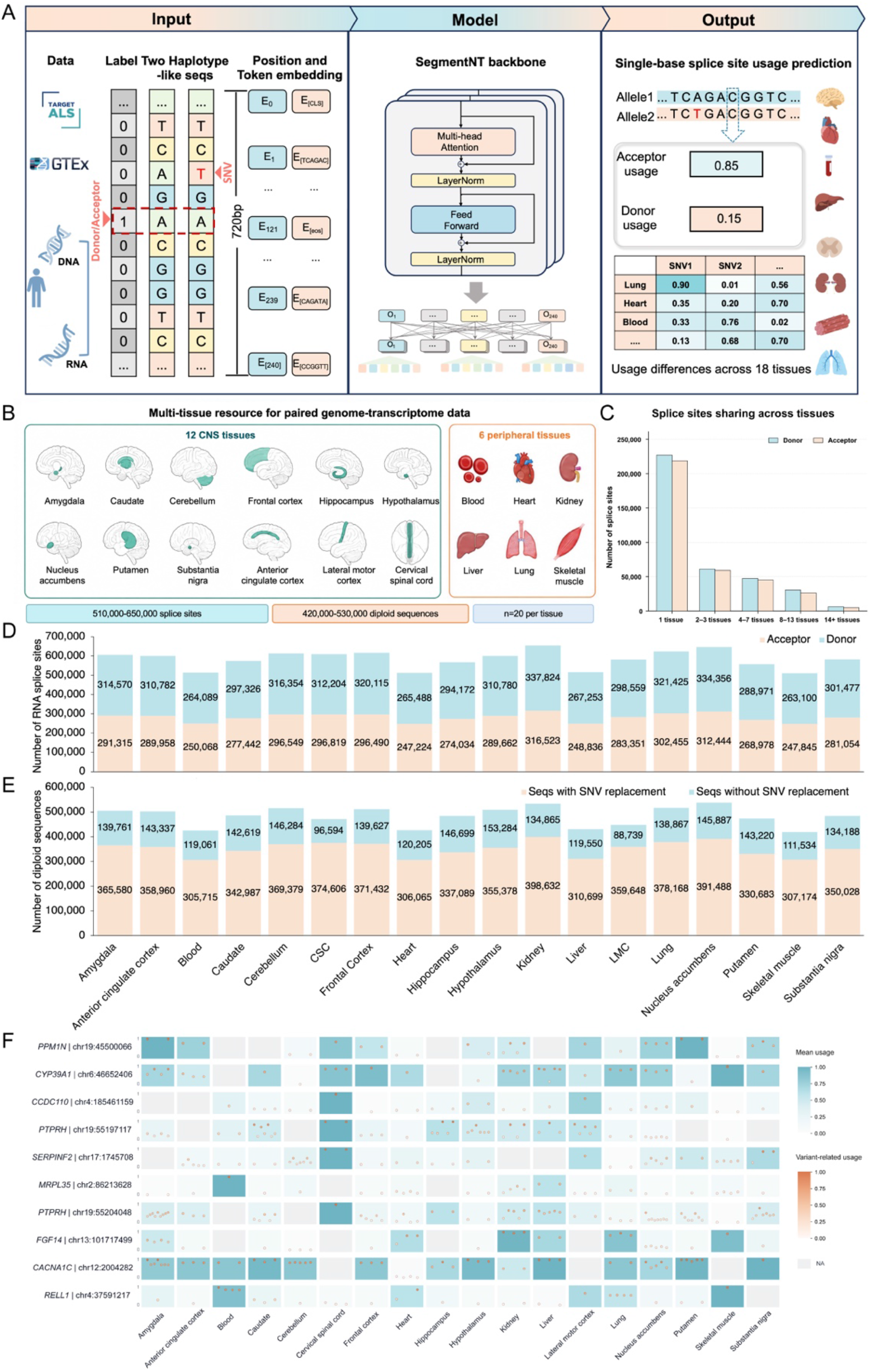
Multi-tissue RNA splice map and Spliformer-V2 architecture. (**A**) Architecture of the Spliformer-V2 model. Spliformer-V2 takes the haplotype-like sequence as input and predicts donor and acceptor site usage scores across human tissues. (**B**) The multi-tissue resource of paired genome and transcriptome data source of 12 human CNS tissues and 6 peripheral tissues derived from TargetALS and GTEx dataset (n=20 per tissue). (**C**) Number of splice sites shared by one or more tissues in the dataset. (**D**) Number of donor and acceptor splice sites in the dataset; (**E**) Number of diploid sequences with/without SNVs replacement in the dataset; (**F**) Heatmap showing representative splice sites with large across-tissue splice usage diversity. The splice sites were selected from those shared in at least 14 human tissues. Splice-sites were ranked by maximum minus minimum of tissue mean usage. Tile color indicates tissue mean usage; gold points indicate variant-related usages positioned on the 0–1 mini-axis.

To model variant effects, we used diploid sequences by substituting individual single nucleotide variants (SNVs) into reference sequence and generating haplotype-like sequences (**Fig. 1A, Fig. S1**). The model used a 720-bp input window and calculated training loss over the central 501 bp, allowing it to capture variant effects within approximately ±250 bp of splice sites^17^, while reducing ambiguity from multiple nearby heterozygous or homozygous variants during haplotype-like sequence reconstruction.

For model development, we constructed a diploid sequence resolved RNA splice map from paired whole genome sequence (WGS) and RNA-seq data across 12 human CNS regions and 6 peripheral tissues from TargetALS and GTEx (n=20 per tissue) (**Fig. 1B**). The RNA splice map contained approximately 510,000–650,000 splice sites and 420,000–537,000 diploid sequences per tissue (**Fig. 1D, E**). Cross-tissue splice-site usage showed a bimodal distribution, with most sites exhibiting either very low or high usage (**Fig.S2**). Most splice sites were detected in one tissue (∼61.3%), whereas only a small fraction (∼1.5%) was shared by at least 14 tissues (**Fig. 1C**). In contrast, the expression of a large portion of protein-coding and noncoding genes was detected across all 18 tissues (**Fig. S3**). Shared splice sites nevertheless showed high across-tissue (**Fig. 1F, S4**) and moderate within-tissue usage diversity (**Fig. S5**), highlighting tissue-specific and variant-related splice-altering effects that are not explicitly modeled by most existing splice-prediction frameworks. Notably > 37.1% splice sites showed across-tissue usage diversity of ≥ 0.25 (P90 − P10) (**Fig. S4**), and > 14.3% splice-tissue pairs exhibited within-tissue sequence-associated usage diversity of ≥ 0.25 (P90 − P10) (**Fig. S5**).

Spliformer-V2 was trained as a regression model to predict splice-site usage scores across tissues. Each tissue-model contains approximately 559 million parameters. Per tissue, the training set included ∼480,000–620,000 splice sites and ∼400,000–510,000 DNA sequences with ∼396,000– 525,000 substituted heterozygous and homozygous variants (**Fig. S6A–C**), whereas the test set included ∼23,000–29,000 splice sites and ∼19,000–26,000 DNA sequences (**Fig. S7A–C**). Approximately 70% of training sequences contained at least one substituted SNV, providing a genotype-informed perturbation landscape for learning variant-associated splicing regulation (**Fig. S6**).

### Performance of Spliformer-V2 and benchmarking

We evaluated Spliformer-V2 on held-out test sets and benchmarked its performance against SpliceTransformer ^14^, Pangolin ^13^, and AlphaGenome ^48^. Spliformer-V2 consistently achieved lower prediction error than other models, with RMSE values of 0.009-0.011, compared with 0.013–0.017 for SpliceTransformer, 0.016–0.022 for Pangolin and 0.018–0.024 for AlphaGenome (**Fig. 2A**). It also showed stronger regression performance, with R² = 0.78–0.86 and Pearson’s r = 0.77–0.86, exceeding SpliceTransformer (R^2^=0.46–0.76, Pearson’s r =0.64–0.79), Pangolin (R^2^=0.53–0.80, Pearson’s r =0.64–0.78) and AlphaGenome (R^2^=0.51–0.73, Pearson’s r =0.71– 0.86) across tissues (**Fig. 2B**).

**Fig. 2.**
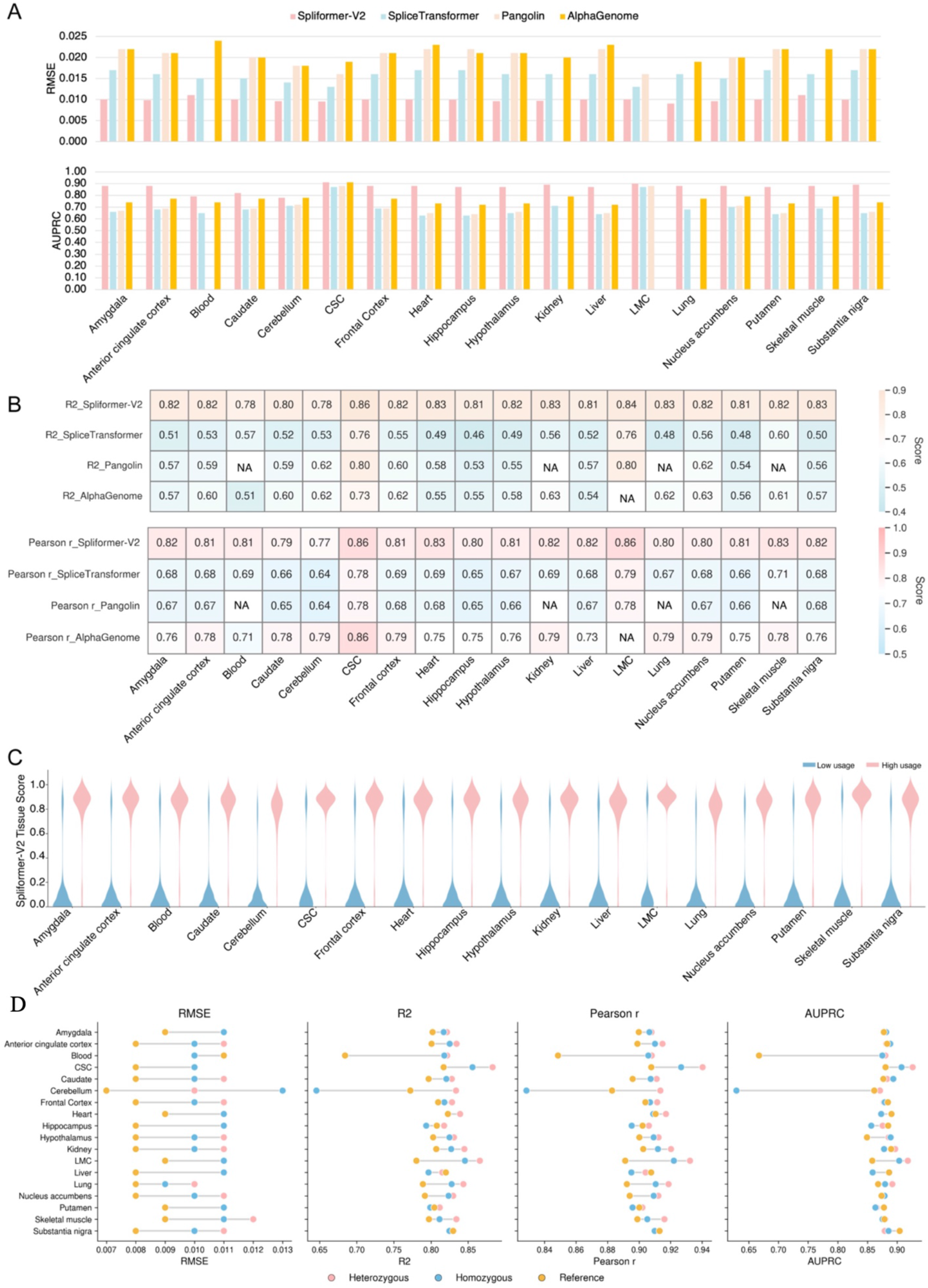
Spliformer-V2 model performance and benchmarking across 18 human tissues. (**A**) RMSE and AUPRC of Spliformer-V2, SpliceTransformer, Pangolin, and Alphagenome on the test set. (**B**) Heatmap showing the R² and Pearson correlation coefficient (r) of the four models on the test set. (**C**) Distribution of Spliformer-V2 predicted splice-site usage scores across 18 tissues. Measured splice sites were classified into low-usage (<0.5) and high-usage (≥0.5) groups. (**D**) RMSE, R², Pearson correlation coefficient (r), and AUPRC of Spliformer-V2 prediction in heterozygous or homozygous SNVs replaced sequences, and reference sequences.

For classification of splice-site usage, Spliformer-V2 achieved higher AUPRC values (0.78–0.91) than SpliceTransformer (0.63–0.87), Pangolin (0.64–0.88) or AlphaGenome (0.72–0.91) across tissues (**Fig. 2A**). Predicted splice-site usage closely matched RNA-seq derived splice usage across 18 tissues (**Fig. 2C**). Notably, Spliformer-V2 showed slightly better performance on heterozygous/homozygous variants-containing sequences than on reference sequences in most tissues, suggesting that the genotype-informed sequence inputs help capture variant-dependent effects on splice-site usage (**Fig. 2D**).

### Performance in identifying rare splicing variants and sQTLs

To assess whether Spliformer-V2 can identify rare variants underlying tissue-specific mis-splicing, we performed outlier-based analysis of paired WGS and RNA-seq data from TargetALS (**Fig. 3A**). In 71-87 aberrant splicing events in lateral motor cortex (LMC) and cervical spinal cord (CSC), Spliformer-V2 recovered 73–81% of splicing outlier-associated variants (Δscore > 0), and 42–56% at Δscore > 0.1 (**Fig. 3B**), indicating its high sensitivity for detecting rare splice-disrupting variants. For example, Spliformer-V2 nominated an *OPTN* variant, chr10:13110032 G>T (hg38), predicted to increase upstream acceptor usage and extend exon 3 in LMC, which was confirmed by RNA-seq (**Fig. 3C**). FRASER analysis showed that this mis-splicing event was absent in other CNS tissues (**Fig. S8**), supporting a tissue-specific effect. In CSC, a *VPS13C* variant, chr15:62044256 C>T (hg38), was predicted to reduce canonical acceptor usage, resulting in exon 2 truncation and a premature termination codon (PTC) (**Fig. 3D**). These examples demonstrate the ability of Spliformer-V2 to prioritize rare splice-altering variants at multi-tissue resolution.

**Fig. 3.**
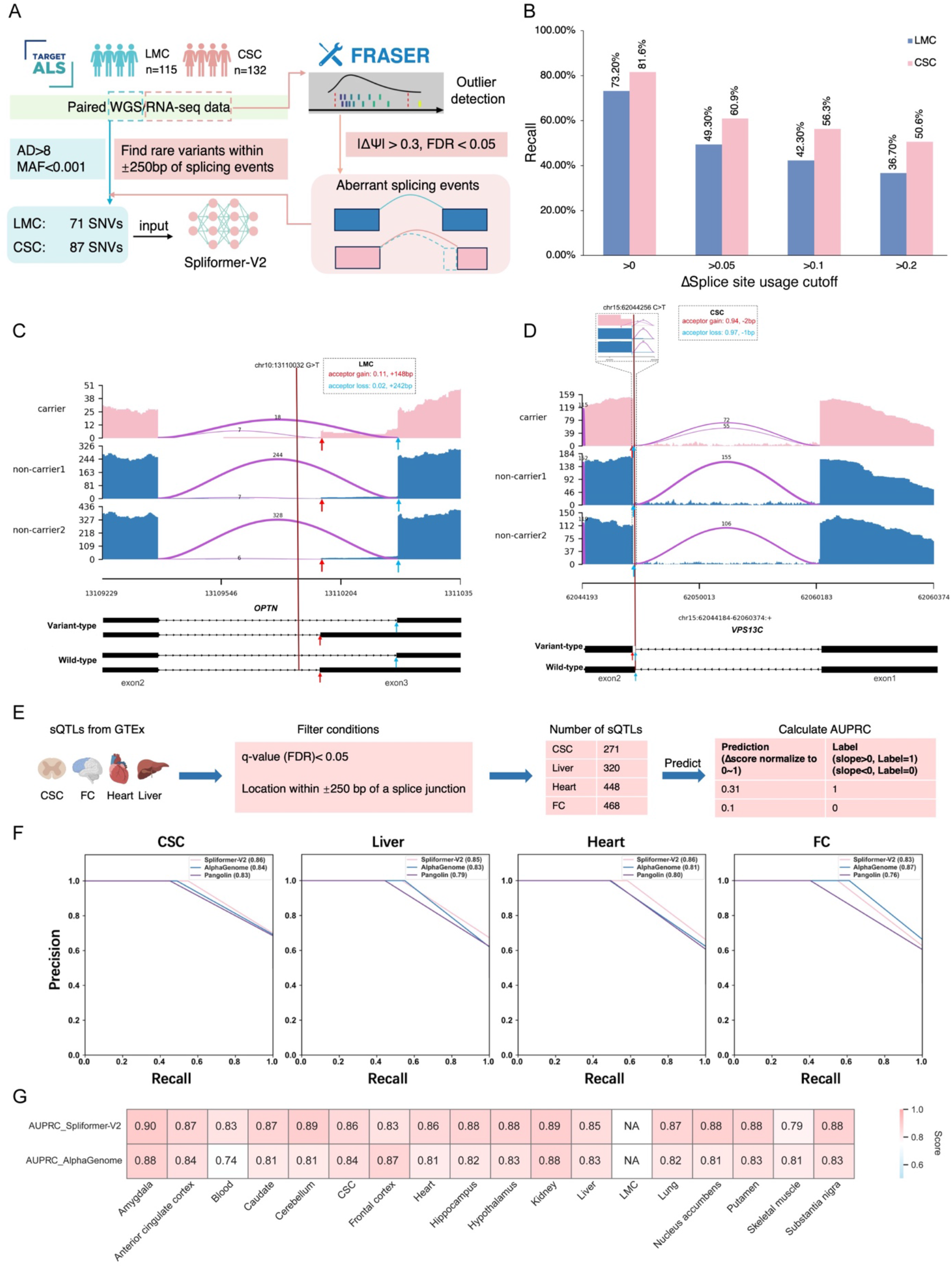
Outlier-based and sQTLs evaluation of Spliformer-V2 model. (**A**) Evaluation workflow. RNA-seq data from the TargetALS database (n=115, LMC; n=132, CSC) were analyzed using FRASER to identify aberrant splicing events. Next, within ±250 bp of the aberrant splicing events, rare variants were assessed in the corresponding samples. In LMC, 71 SNVs were identified to be associated with aberrant splicing events, while 87 SNVs were identified in CSC. Finally, Spliformer-V2 was used to predict whether these SNVs had splicing effects consistent with the aberrant splicing events observed. (**B**) Model recall at different thresholds of splice site usage. (**C-D**) Examples showing the validation of rare splicing variants based on RNA-seq data. A variant in *OPTN* (chr10:13110032 G>T, hg38) increases the usage of the acceptor site at -7 bp from the variant, causing the extension of exon 3 (**C**). A variant in *VPS13C* (chr15:62044256 C>T, hg38) decreases the usage of the acceptor site at -1 bp while introducing a novel acceptor site at -2 bp, leading to truncation of exon 2 and a PTC (**D**). **(E)** Workflow for evaluating model performance on GTEx sQTLs. sQTLs from CSC, FC, heart, and liver were obtained from the GTEx portal and filtered for FDR q-value < 0.05 and localization within ±250 bp of a splice junction. Model-predicted Δscores were normalized to a range of 0-1. Variants with positive and negative sQTL slopes were assigned labels of 1 and 0, respectively, and AUPRC was calculated to assess prediction performance. **(F-G)** Performance of Spliformer-V2, Pangolin and AlphaGenome in predicting the directional splicing effects of GTEx sQTLs across human tissues: (**F**) AUPRC curves showing the performance of models for classification task in 4 tissues; (**G**) Heatmap showing AUPRC values of AlphaGenome and Spliformer-V2 in 18 tissues.

We next evaluated Spliformer-V2 on the direction of splicing quantitative trait loci (sQTLs). Spliformer-V2 outperformed Pangolin in classifying alerted splice-site usage in CSC, frontal cortex (FC), heart and liver (**Fig. 3E, F**). Additionally, across 15 of 17 tissues, Splifomer-V2 had better performance than AlphaGenome for evaluating sQTLs (**Fig. 3G**).

### Interpreting tissue-specific splicing motifs

To assess whether Spliformer-V2 captures tissue-specific splicing regulatory grammar, we analyzed the Ascot dataset and identified 657 differential cryptic exon (CE) events between CSC and blood (|ΔPSI| > 0.3). Integrating these events with Spliformer-V2 predictions prioritized 100 CEs with concordant acceptor and donor changes (|Δscore| > 0.1), including 58 CEs with higher PSI in CSC (**Fig. 4A**). In silico mutagenesis (ISM) analysis identified sequence features contributing to tissue-specific CE usage. In blood, ISM signals around donor and acceptor sites were generally weak, with only modest changes near canonical splice sites. By contrast, CSC showed stronger signals at canonical donor GT and acceptor AG dinucleotides, as well as in the upstream region of acceptor (−20 to -5 bp), consistent with the polypyrimidine tract (**Fig. 4B, C**).

**Fig. 4.**
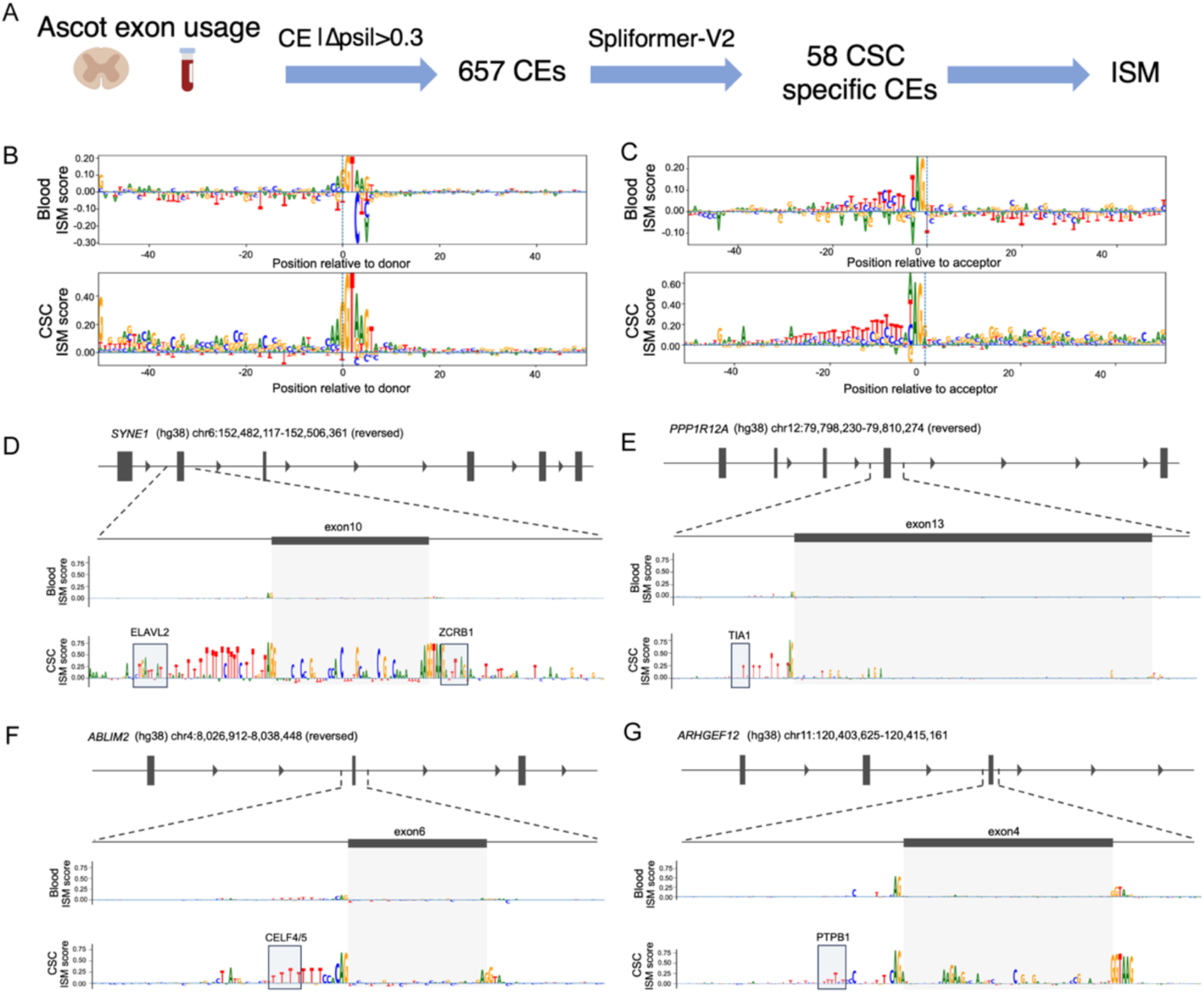
CSC-specific splicing events and RBP-associated regulatory motifs. **(A)** Workflow for identifying CSC specific CE events and analyzing their regulatory motifs using ISM. (**B-C**) ISM profiles centered on splice donor (**B**) and acceptor (**C**) sites in blood (top) and CSC (bottom). The x-axis indicates position relative to the splice site (0 denotes the splice site), and the y-axis represents ISM score. **(D–G)** Example loci in *SYNE1* (**D**), *PPP1R12A* (**E**), *ABLIM2* (**F**), and *ARHGEF12* (**G**) showing CSC-specific splicing regulatory features. The top panel displays gene structure, with the analyzed exon shown in a zoomed-in view below. ISM profiles reveal minimal signal in blood, whereas CSC exhibits pronounced splicing regulatory activity across exonic and flanking intronic regions, indicating stronger sequence-dependent regulation. Motif analysis identifies enrichment of RBPs, including ELAVL2, TIA1, CELF4/5, and PTBP1.

Motif analysis indicated high-contribution regions overlapped known RNA-binding protein (RBP) motifs, including neuron-related RBPs such as ELAVL2 and CELF4/5^18, 19^, and general splicing factors including TIA1, U2AF2, PTBP1, HNRNPK and ZCRB1 (**Table S1**). For example, ISM analysis identified an ELAVL2 motif approximately 42 bp to *SYNE1* splice-site (**Fig. 4D**), and a TIA1 motif (TTTTT) close to *PPP1R12A* splice-site (**Fig. 4E**). Furthermore, CELF4/5 and PTBP1 motifs are close to CSC-specific *ABLIM2* and *ARHGEF12* cryptic exons (**Fig. 4F, G**). Together, these findings show that Spliformer-V2 can reveal tissue-specific splicing motifs and nominate candidate RBPs underlying CNS-specific splice regulation.

### Tissue vulnerability of disease-associated splice-altering variants

Since splice-altering variants are strongly linked to pathogenicity, we first assessed their tissue-specific enrichment using 1,256 pathogenic and 105,171 benign variants from ClinVar (**Fig. S9, Table S2**). Across 18 tissues, pathogenic variants showed a significantly higher fraction of predicted splice-altering variants than benign variants (Fisher’s exact test, P < 2.2 × 10⁻¹⁶) (**Fig. S9**). At |Δscore| > 0.9, LMC showed the highest fraction of pathogenic splice-altering variants among CNS tissues (24.2% versus 3.7–12.7%), whereas skeletal muscle showed the highest fraction among peripheral tissues (24.8% versus 4.6–11.3%) (**Fig. S9**). These findings suggest increased vulnerability of LMC and skeletal muscle to pathogenic splice disruption.

We next examined tissue vulnerability in neurodegenerative disease by analyzing rare variants across 60 ALS-associated genes across 1,405 ALS samples with WGS data (**Fig. S10A, Table S3**). Splice-altering variants harbored in ALS-associated genes were enriched in LMC and skeletal muscle relative to other tissues (**Fig. S10B**). At the gene level, *PTPRN2*, *UNC13A* and *CHMP2B* showed higher fractions of predicted splice-altering variants in LMC and skeletal muscle, whereas *SCFD1* showed a higher fraction in FC (**Fig. S10C–F**). Finally, using Z-score distributions to quantify splicing variant effect size^14^, we observed a greater burden of large-effect splice-altering variants in CSC and LMC than in FC (CSC, 92%; LMC, 64%; FC, 44%) (**Fig. S10G–I**). Together, these analyses indicate that Spliformer-V2 can resolve tissue-specific vulnerability to disease-associated splice-altering variation, highlighting a mechanism potentially underlying susceptible motor neuron-relevant tissues in ALS.

### CNS tissue-specific mis-splice vulnerable genes in neurological diseases

To fine-map disease-associated noncoding variants with splice-altering effects across tissues, we first analyzed 2,158 ALS-associated variants from a large GWAS meta-analysis ^20^. Spliformer-V2 identified 77 splice-altering variants, of which 33 (43%) showed CNS-specific effects (**Fig. S11A, B, Supplementary File 1**). These included multiple splice-altering variants at *MOB3B-C9ORF72* locus (rs12554036, rs10738774, rs1537712 and rs2492818), the *SCFD1* locus (rs10142331, rs2273521, rs230366 and rs9322864) (**Fig. S11C, D**), and the previously implicated splice-vulnerable gene *UNC13A* (rs78549703)^9, 10^. Brain sQTL-based SMR-portal analyses^21^ further supported the links between genetically regulated splicing of *C9ORF72*/*SCFD1* and ALS risk, suggesting that *C9ORF72-MOB3B* and *SCFD1* represent CNS-specific mis-splice vulnerable genes contributing to ALS susceptibility.

Furthermore, Spliformer-V2 analyses of neurodegenerative disease GWAS data ^22, 23^ identified 10 AD-, 124 PD- and 14 frontotemporal dementia (FTD)-associated splice-altering variants, of which 64.2–90% showed CNS-specific effects (**Fig. S12A–C, Supplementary File 2**). Notable variants mapped near *SNCA* (rs78504001), *APOE* (rs769449) and *BTNL2* (rs3129953), with SMR-portal analyses supporting brain sQTL links for *SNCA* in PD and *APOE* in AD.

Finally, analyses of neuropsychiatric GWAS loci^22^ identified 11 autism spectrum disorder-, 21 bipolar disorder-, 5 schizophrenia- and 2 depression-associated splice-altering variants (**Fig. S13A–D, Supplementary File 3**), including candidate variants in *PRPF19* (rs140741951), *IGLL1* (rs115907934), *CCSER1* (rs7685994) and *GCLC* (rs80319574 and rs17882530). Together, these results show that Spliformer-V2 can prioritize CNS-specific splice-altering variants and identify mis-splice vulnerable genes in neurological diseases.

### Prediction of rare CSC/LMC splice-altering variants nominates *PTPRN2* in ALS

To assess the contribution of rare splice-altering variants to ALS, we analyzed rare intronic variants from 1,405 ALS WGS samples (MAF < 0.0001, Δscore ≥ 0.4; **Table S3**). Spliformer-V2 nominated ∼11,000–16,000 rare splice-altering variants in CSC or LMC, including 46–77 splice-altering variants on ALS genes (**Fig. 5, S14**), highlighting *PTPRN2* and *ERBB4*. Notably, 7 or 6 patients harbored extremely rare *PTPRN2* variants (absent from the gnomAD v3.12 database) predicted to be splice-altering in CSC or LMC, respectively, supporting its potential contribution to ALS risk. Minigene assays in SH-SY5Y cells confirmed that the *ERBB4* variant chr2:211919854 A>T (hg38) generated an 89-bp cryptic exon in intron 3 with a downstream PTC (**Fig. S15A–C**). Similarly, the *PTPRN2* variant rs555566959, chr7:158122466 C>T (hg38), generated a 100-bp cryptic exon in intron 9, also introducing a PTC (**Fig. S15D–F**).

**Fig. 5.**
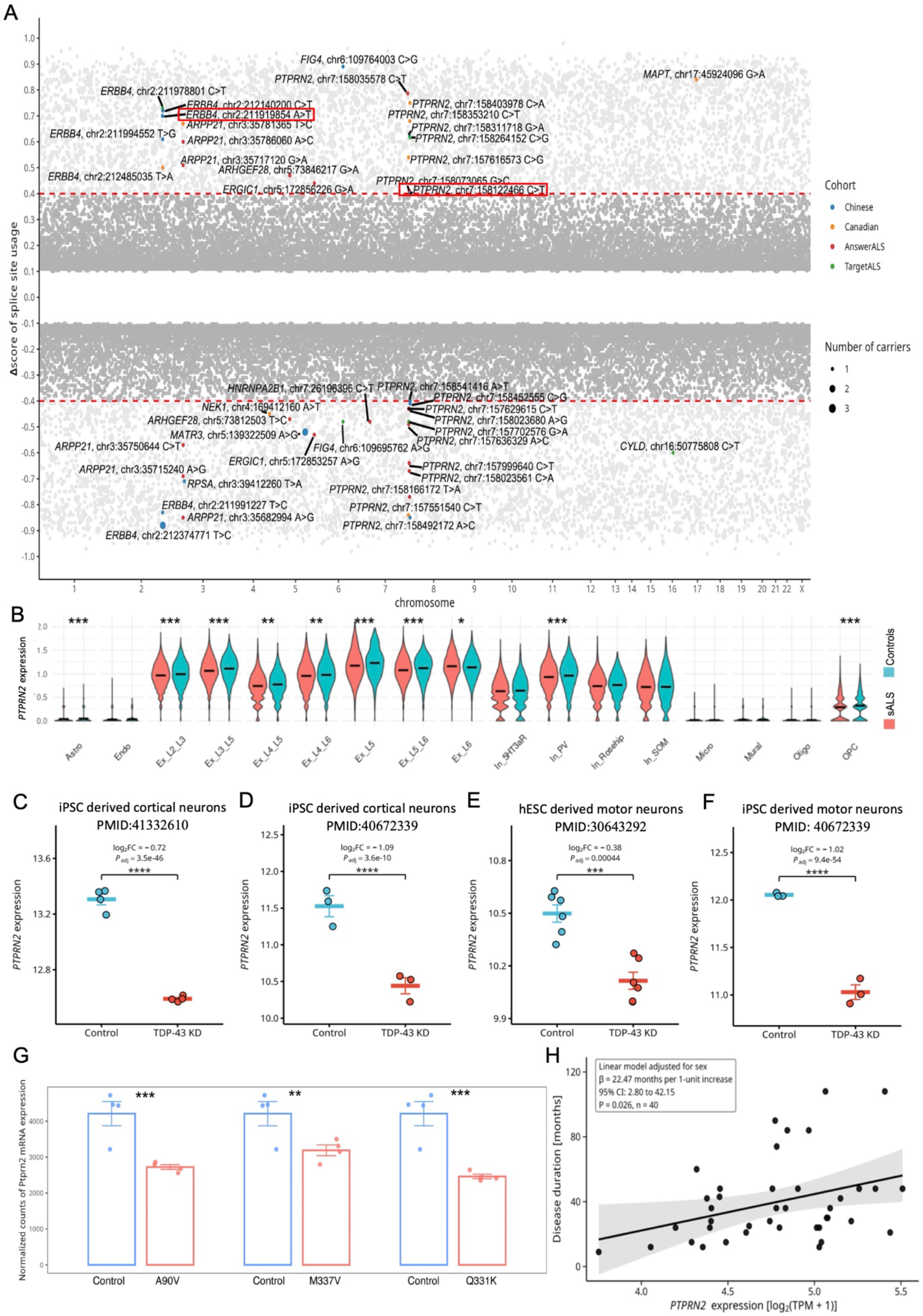
Rare splicing variants in ALS WGS and altered *PTPRN2* expression in ALS disease or pathology associated neurons. **(A)** The Dot plot showing predicted CSC rare splicing variants in ALS WGS (n=1,405). The x-axis represents chromosomes, and the y-axis represents Δscore predicted by Spliformer-V2. Point size corresponds to carrier numbers. ALS genes were labeled. The red rectangle indicated rare variants selected for minigene assays. (**B**)Violin plots showing the differential expression of *PTPRN2* at single-cell resolution in motor cortex (17 sALS and 16 controls, Synapse: syn51105515). Two rows correspond to one gene and two columns represent a cell type, with red representing the ALS patients and blue representing the controls. The y-axis represents the log10-converted count values. Significance was determined by Wilcoxon rank-sum test (Benjamini-Hochberg adjusted p value): *** P.adj < 0.001; ** P.adj < 0.01; * P.adj < 0.05. **(C-F)** Re-analysis of published RNA-seq datasets in neurons showed reduced *PTPRN2* expression upon TDP-43 knockdown in iPSC-derived cortical neurons (**C**, log₂FC = -0.72, adjusted *P* = 3.5 × 10^-^⁴⁶, n = 4 per group; **D**, log₂FC = -1.09, adjusted *P* = 3.6 × 10^-^¹⁰, n = 3 per group), hESC-derived motor neurons (**E**, log₂FC = -0.38, adjusted *P* = 4.4 × 10^-^⁴, n = 6 per group), iPSC-derived motor neurons (**F**, log₂FC = -1.02, adjusted *P* = 9.4 × 10^-^⁵⁴, n = 3 per group). Each point represents an independent biological replicate. Data are presented as mean ± SEM. Expression values are shown as log₂(DESeq2-normalized counts + 1). Differential expression was assessed using the DESeq2 Wald test, and *P* values were adjusted for multiple testing using the Benjamini-Hochberg method. ***adjusted *P* < 0.001; ****adjusted *P* < 0.0001. **(G)** Bar figures showing the *Ptprn2* mRNA expression in rat neurons expression different TDP-43 mutants. Quantifications are presented as mean ± SEM. P-values were calculated using the Wald test and adjusted using the Benjamini-Hochberg method. Significance levels are indicated as follows: *** P < 0.001; ** P < 0.01; * P < 0.05. **(H)** Scatter plot showing the relationship between *PTPRN2* expression and ALS disease duration in TargetALS thoracic spinal cord RNA-seq. Each point represents one independent ALS donor (n = 40). *PTPRN2* expression is shown as log₂(TPM + 1), and disease duration is shown in months. The solid line represents the sex-adjusted marginal fitted association from a multivariable linear regression model, and the shaded area indicates the 95% confidence interval of the fitted line.

Furthermore, ISM analysis showed that 6 *PTPRN2* splice-altering variants may create splice sites directly, and 15 may alter RBP-binding motifs (**Table S4**), therefore increasing ISM scores. For example, the rs555566959 variant introduced a novel splice acceptor (**Fig. S16**), whereas other variants altered binding motifs for RBPs including SRSF5/9, ELAVL2/TIA1, SRSF1/SRSF9, GRSF1/HNRNPF, PTBP1 and SRSF6 (**Fig. S17A–F**). These results show that *PTPRN2* rare variants may modify splice motifs or RBP binding motifs.

*PTPRN2* is a marker of L5 and L3/L5 excitatory neurons (**Table S5**), vulnerable cell types in ALS ^12, 24^. Analysis of human motor cortex snRNA-seq data ^24^ showed that *PTPRN2* was significantly downregulated in multiple excitatory neuronal subtypes in sporadic ALS compared to controls, including Ex_L5 and Ex_L3_L5 neurons (**Fig. 5B**). Consistently, TDP-43 knockdown in iPSC or hESC derived neurons ^8, 25, 26^ reduced *PTPRN2* mRNA expression by approximately 23–53% (**Fig. 5C–F**) that were often accompanied by multiple cryptic splicing events (**Fig. S18**). *Ptprn2* was also downregulated in rat hippocampal neurons expressing ALS disease-causing TDP-43 mutants (A90V, M337V and Q331K) (**Fig. 5G**). Notably, higher *PTPRN2* expression in thoracic spinal cord was associated with longer disease duration (**Fig. 5H**), consistent with a recent report linking higher CSF PTPRN2 protein levels to slower ALS progression ^27^. Together, these findings nominate *PTPRN2* as a mis-splice vulnerable ALS gene and suggest that reduced neuronal *PTPRN*2 expression underlies ALS/TDP-43 pathology.

### *PTPRN2* overexpression rescues *C9ORF72* ALS/FTD iMNs degeneration and modulates TDP-43 mislocalization

Two independent studies reported reduced CSF PTPRN2 levels in ALS and FTD patients^28, 29^. We found that TDP-43 depletion in NGN2-induced human neurons (iNs) reduced overall *PTPRN2* transcript levels (**Fig. 6A–B**), consistent with observations in postmortem ALS and FTD cortex, and *in vitro* studies describing increased *PTPRN2* intronic cryptic splicing and lower *PTPRN2* mRNA levels after TDP-43 knockdown ^30^. In addition, TDP-43 knockdown (∼50%) reduced PTPRN2 protein levels by approximately 20% (**Fig. 6C–E**).

**Fig. 6.**
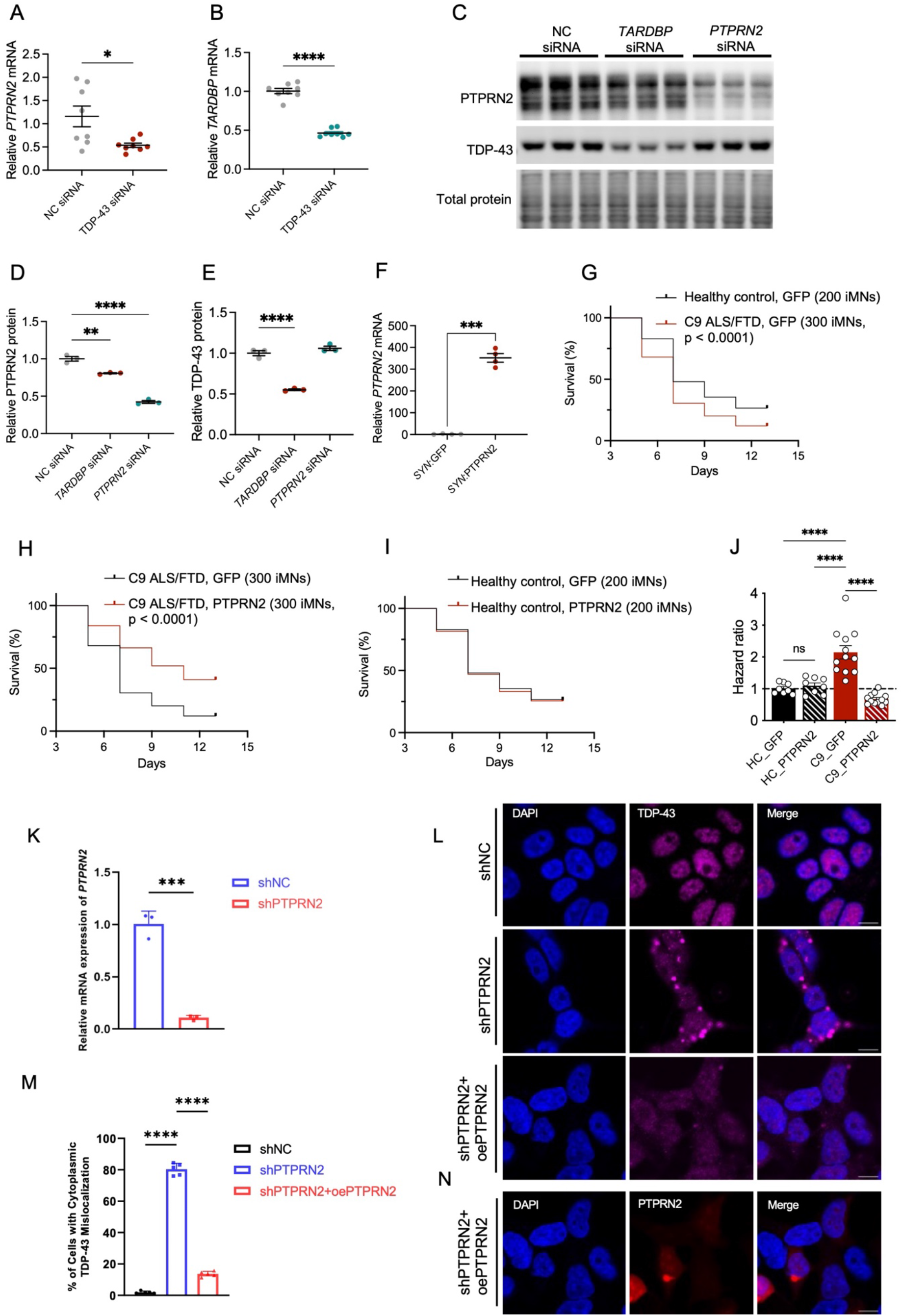
PTPRN2 overexpression rescues degeneration of *C9ORF72* ALS/FTD iMN and modulates TDP-43 mislocalization. **(A-J)** *PTPRN2* rescues iMN survival derived from *C9ORF72* ALS/FTD patients. (**A, B**) *PTPRN2* mRNA expression was significantly reduced following TDP-43 knockdown in induced neurons (iNs). n = 8 per condition; t-test; *p < 0.05, ****p < 0.0001. (**C – E**) Western blot analysis demonstrated reduced PTPRN2 protein levels following TDP-43 knockdown in iNs. n = 3 for each condition; one-way ANOVA; **p < 0.01, ****p < 0.0001. (**F**) Lentiviral overexpression of *PTPRN2* driven by the SYN1 promoter (SYN1:PTPRN2) significantly increased *PTPRN2* mRNA expression in iNs compared with the SYN1:GFP control. n = 4; t-test; ***p < 0.001. (**G – I**) Kaplan–Meier survival analysis of iMNs derived from healthy controls (two lines per condition) and *C9ORF72* ALS/FTD patients (three lines per condition) following transduction with either SYN1:GFP or SYN1:PTPRN2:GFP. Survival curves were compared using the logrank (Mantel–Cox) test. (**J**) Hazard ratios for G – I, calculated using Mantel-Haenszel method. Bar graphs represent means ± SE, one-way ANOVA; ****p < 0.0001. (**K-N**) PTPRN2 modulates TDP-43 mislocalization. **(K)** Relative *PTPRN2* mRNA expression (adjusted to *GAPDH*) in SH-SY5Y cells with RP-qPCR upon *PTPRN2* knockdown. Cells were transfected with shNC or shPTPRN2. Data are presented as mean ± SEM (n = 3 replicates). ***p < 0.001 vs. shNC (unpaired two-tailed Student’s t-test). **(L)** Representative immunofluorescence images showing TDP-43 subcellular localization. Cells were transfected with shNC, shPTPRN2, or co-transfected with shPTPRN2 and the *PTPRN2* overexpression plasmid (shPTPRN2+oePTPRN2). The magenta color represents TDP-43, the blue color represents nuclei with DAPI. Scale bar = 10 μm. **(M)** Quantification of the percentage of cells exhibiting abnormal cytoplasmic TDP-43 aggregation. Data are presented as mean ± SEM from 5 randomly selected fields of view (FOV) per condition. ****p < 0.0001 for the indicated comparisons (one-way ANOVA followed by Tukey’s multiple comparisons test). **(N)** Representative immunofluorescence images showing the expression of the *PTPRN2* plasmid in SH-SY5Y cells. The mCherry fluorescent tag (red) indicates successful overexpression, with nuclei counterstained by DAPI (blue). Scale bar = 10 μm.

To test whether PTPRN2 enhances neuron survival in disease-relevant contexts, we overexpressed *PTPRN2* in induced motor neurons (iMNs) generated from two healthy controls and three *C9orf72* ALS/FTD patient-derived iPSC lines. As we previously reported, *C9ORF72* ALS/FTD iMNs degenerated significantly faster than controls upon withdrawal of neurotrophic factors from the media (**Fig. 6G, S19A**). qRT-PCR confirmed robust induction of *PTPRN2* after transduction with *SYN1*::PTPRN2-GFP (**Fig. 6F**). *PTPRN2* overexpression significantly improved the survival of *C9ORF72* FTD/ALS iMNs (**Fig. 6H, J, S19B**), rescuing their survival to the level of control iMNs **(Fig. 6G, J, S19A**). In contrast, *PTPRN2* overexpression did not alter survival of control iMNs (**Fig. 6I, J, S19C**), suggesting a disease-context-specific protective effect rather than a general pro-survival effect.

We then examined whether *PTPRN2* influences TDP-43 cytoplasmic mislocalization, a hallmark of ALS/FTD pathology. In SH-SY5Y cells, *PTPRN2* knockdown markedly increased cytoplasmic TDP-43 mislocalization (p < 0.0001, fold change = 40.7; **Fig. 6K, L**), whereas *PTPRN2* overexpression partially rescued TDP-43 mislocalization (p < 0.0001, fold change = 5.9; **Fig. 6M, N**). Together, these results suggest that *PTPRN2* restoration could promote neuronal resilience in *C9ORF72* ALS/FTD iMNs, potentially by mitigating TDP-43 proteinopathy, and nominate *PTPRN2* as a therapeutic modifier in ALS.

## Discussion

Dissecting tissue or cell specific splicing regulation is essential for understanding transcriptomic diversity and disease mechanisms ^2, 31, 32^. Here, we generated a multi-tissue, diploid sequence-resolved RNA splice map and developed Spliformer-V2 to predict splice-site usage across 12 human CNS regions and 6 peripheral tissues. Spliformer-V2 enabled accurate multi-tissue prediction, interpretation of regulatory motifs and prioritization of disease-associated splice-altering variants. Application to neurological disease loci identified CNS-specific mis-splice-vulnerable genes. In ALS WGS data, Spliformer-V2 nominated rare splice-altering variants enriched in *PTPRN2*, which showed reduced neuronal expression under TDP-43 loss-of-function. Functionally, *PTPRN2* overexpression rescued survival of *C9ORF72* ALS/FTD iMNs and mitigated TDP-43 cytoplasmic mislocalization.

Our unique RNA splice map spanned approximately 420,000–537,000 genotype-informed diploid sequences and 510,000–650,000 splice sites across human tissues, including unreported CNS regions like LMC that are highly relevant to neurodegenerative disease. This resource complements existing GTEx-based sQTL^4^ and SpliceMap ^17^ datasets by linking individual genetic variation to tissue-resolved splice-site usage.

By incorporating genotype-informed diploid sequence, Spliformer-V2 can distinguish heterozygous and homozygous variant effects and partially capture compound effects of nearby variants ^33^. Built on pretrained nucleotide representations learned from billions of genomic tokens^16^, Spliformer-V2 transfers general sequence grammar to tissue-specific splicing prediction. It provides single-nucleotide-resolution output that enables ISM analysis, allowing regulatory features such as canonical splice-site elements and RBP-binding motifs to be interpreted in a tissue-specific manner. That is a key advance compared to statistical models for common and rare splice-altering variants ^4^ ^17^ .

Brain is among the most splicing-complex tissues ^34^, yet many existing models^13, 14^ treat brain as a single mixed tissue. By resolving multiple CNS subregions, Spliformer-V2 revealed tissue-specific vulnerability to splice-altering variants. LMC and skeletal muscle showed particularly high burdens of predicted pathogenic splice-altering variants, indicating the idea that tissue-specific splicing regulation contributes to selective vulnerability in disease ^35^.

ALS provided a disease context for translating variant prediction into biological insight. TDP-43 pathology causes widespread RNA mis-splicing in ALS/FTD^7–10, 26, 36, 37^, and many ALS genes harbored RNA splice-altering variants^12^. Analysis of ALS genomes prioritized rare splice-altering variants enriched in *PTPRN2* with predicted loss-of-function effects. Notably, *PTPRN2* is a TDP-43 bound transcript ^38, 39^, and is consistently downregulated in TDP-43-depleted iPSC derived neurons^8, 25, 26^. Its overexpression rescued *C9ORF72* ALS/FTD iMNs degeneration and reduced TDP-43 mislocalization, supporting *PTPRN2* as a potential therapeutic target for AAV-mediated gene delivery or ASO-based ^40^ splice-correction intervention.

Several limitations remain. First, the current model uses genotype-informed sequences rather than fully phased haplotypes. Statistical phasing methods such as SHAPEIT^41^ or Beagle ^42^ could improve haplotype reconstruction, although rare variants remain difficult to phase accurately. Second, the splice map is based on short-read RNA-seq, which may miss complex isoforms and long transcript structures abundant in brain. Single-cell long-read sequencing could further improve splicing annotation ^43^. Third, tissue-specific prediction may benefit from explicitly incorporating RBP expression or activity ^44^, which could help model both cis-/trans-regulatory code across tissues. Finally, tissue specificity may reflect differences in the magnitude of splice change; thus, cell-line RNA alterations may resemble those in motor neurons to varying degrees.

In summary, Spliformer-V2 provides a framework for accurate and interpretable prediction of splice-altering variants across human tissues. Its application to neurological disease uncovers tissue-specific mis-splice vulnerability and nominates *PTPRN2* as an ALS modifier, highlighting the value of multi-tissue splicing models for variant interpretation and therapeutic target discovery.

## Methods

### Data preparation

To generate a diploid sequence resolved RNA splice map, we used paired WGS and RNA-seq data from TargetALS and GTEx (**Table S6**), including 12 CNS regions and 6 peripheral tissues. GTEx tissues included amygdala, anterior cingulate cortex, caudate, cerebellum, frontal cortex, hippocampus, hypothalamus, nucleus accumbens, putamen, substantia nigra, blood, heart, kidney, liver, lung and skeletal muscle. TargetALS provided LMC and CSC data. For each tissue, we included 10 ALS patients and 10 controls from TargetALS or 20 controls from GTEx.

GTEx RNA-seq reads were aligned to the human reference genome hg38 using STAR v2.7.9a with *--twopassMode Basic*. For TargetALS, pre-aligned BAM files were used. Splice junctions were extracted using RegTools v0.5.2, and splice-site usage was quantified using SpliSER v1.3. Splice sites supported by fewer than five reads were excluded. Sites with maximum usage below 0.05 across all individuals were relabeled as non-splice sites.

To construct diploid sequence inputs, we first extracted 720-bp reference sequences from hg38 using bedtools v2.30.0. Genomic coordinates were derived from the longest protein-coding transcript for each gene in GENCODE v39. Sequences were then generated using two strategies. First, splice site-centered sequences were constructed by extracting 720-bp regions centered on each splice site. Second, a sliding-window approach was applied from each transcription start site using a window size of 720 bp and a step size of 501 bp. Only windows containing at least one splice site within their central 501 bp were retained. Sequences from both strategies were merged to generate the final dataset. SNVs were incorporated from WGS VCF files, retaining variants with allele depth greater than 8.

To model genotype-related splicing effects, SNVs were substituted into the reference sequence under three conditions: one heterozygous variant; one or more homozygous variants; or one heterozygous variant together with one or more homozygous variants (**Fig. S1**). Because phased haplotypes were not directly available, heterozygous variants were randomly partitioned into two haplotype-like sequences as previously described ^45^. Each heterozygous SNV was introduced into one haplotype sequence while the other allele remained unchanged, and the assignment was then switched to generate the alternative haplotype-like sequences. Homozygous SNVs were introduced into both alleles.

After SNV substitution across all individuals, sequences were merged into tissue-specific datasets. To ensure label consistency, duplicate sequences were retained only when all pairwise differences in splice-site usage were less than 0.05. For retained duplicates, splice-site usage labels were averaged. The two haplotype-like sequences were concatenated into a single diploid sequence input using two special tokens: <CLS> was prepended to the first allele and <EOS> to the second. The full processing workflow is shown in **Fig. S1**.

Diploid sequences from chromosomes 2, 4, 6, 8 and 10–22 were used for training, whereas chromosomes 1, 3, 5, 7 and 9 were used for testing. To minimize data leakage from homologous sequence similarity, paralogous genes were excluded from the test set using the Ensembl BioMart human paralog list.

### Model architecture and training

The architecture of the Spliformer-V2 model is shown in **Fig. 1A**. We initialized Spliformer-V2 with the model weights from SegmentNT multi-species, a pretrained DNA foundation model based on Nucleotide Transformer representations^15, 16^, for fine-tuning. SegmentNT contains an input embedding layer, a BERT-based encoder and an output module.

In the input layer, nucleotide sequences were tokenized into 6-mers using the *AutoTokenizer* module, mapped to integer indices and subjected to a random masking strategy. Each token is embedded through the sum of its token embedding and corresponding positional encoding.

The BERT-based encoder consists of 24 Transformer layers. Each layer includes multi-head self-attention, residual connections, layer normalization and a position-wise feed-forward network with GELU activation. To enable nucleotide-level splice-site prediction, the original masked-language-modeling head was replaced with a one-dimensional U-Net segmentation module. The U-Net module contains two downsampling and two upsampling convolutional blocks with 1,024 and 2,048 filters. A final classification output head predicts donor and acceptor splice-site usage scores.

Separate tissue-specific models were trained on data from each of the 18 tissues. Models were trained for 20 epochs with a batch size of 24 on a single NVIDIA A100 PCIe 40-GB GPU. The initial learning rate was 5 × 10⁻⁵. AdamW was used for optimization, and mean squared error (MSE) was used as the training loss.

### Model performance evaluation and benchmarking

Spliformer-V2 was evaluated on the held-out test set (**Fig. S7**) and benchmarked against SpliceTransformer, Pangolin and AlphaGenome. For Spliformer-V2, 720-bp inputs were used, and evaluation was restricted to the central 501 bp. For SpliceTransformer and Pangolin, 4,000 bp and 5,000 bp flanking sequences, respectively, were added on each side. For AlphaGenome, 5,000 bp flanking sequences were retained on each side and N bases were padded to obtain a total input length of 16,384 bp. Evaluation for all models was restricted to the same central 501-bp region.

Because SpliceTransformer, Pangolin and AlphaGenome accept haploid sequences, heterozygous variants were treated as homozygous when generating individual-specific sequences for these models. For CNS tissues, SpliceTransformer and Pangolin predictions from the general brain model were used because CNS subregion-specific models were unavailable. Pangolin was not benchmarked in blood, kidney, lung or skeletal muscle because corresponding tissue models were unavailable. AlphaGenome was not evaluated in LMC because this tissue type is not represented in the model.

We quantified model performance using root mean squared error (RMSE), coefficient of determination (R²), Pearson correlation and area under the precision–recall curve (AUPRC). For AUPRC, splice-site usage greater than 0.5 was labeled as high usage and all other sites as low usage. Metrics were computed using *average_precision_score, r2_score* and *mean_squared_error* from sklearn.metrics, and Pearson correlation was calculated using *pearsonr* from scipy v1.7.1.

### Rare splice-altering variant evaluation

To evaluate the ability of Spliformer-V2 to identify rare splice-altering variants, we analyzed RNA-seq data from TargetALS in LMC (n = 115) and CSC (n = 132). Aberrant splicing events were identified using FRASER ^46^. For each event, we searched for rare variants within ±250 bp of the affected splice junction^17^. We used Spliformer-V2 to predict the splicing effects of these variants, and the predictions were compared with the FRASER-derived aberrant splicing patterns. Candidate events were visualized using Trackplot v0.5.3.

### sQTLs evaluation

To evaluate prediction of splicing quantitative trait loci (sQTLs), we selected GTEx sQTLs from 4 tissues (CSC, FC, heart and liver) that could be predicted by both Spliformer-V2 and Pangolin, and from 17 of the 18 tissues (except for LMC) that could be predicted by both Spliformer-V2 and AlphaGenome. sQTLs were retained if they had a false discovery rate (FDR) q-value < 0.05 and were located within ±250 bp of a splice junction. Variants with positive effect sizes (slope > 0) were labeled as 1, and those with negative effect sizes (slope < 0) were labeled as 0. Precision-recall curves were generated using *precision_recall_curve*, and AUPRC was calculated using *average_precision_score* from scikit-learn.

### In silico mutagenesis and motif analysis

Tissue-specific cryptic exon events between CSC and blood were identified from the Ascot dataset ^47^. Cryptic exons with |ΔPSI| > 0.3 were retained, yielding 657 events. Spliformer-V2 was applied to both tissues, and events with concordant predicted changes in donor and acceptor usage were retained using |Δscore| > 0.1. This yielded 100 high-confidence events, including 58 cryptic exons with higher usage in CSC. We performed in silico mutagenesis as reported ^48^ to identify high-contribution sequence regions. Candidate motifs were scanned against known motif databases using FIMO ^49^. For analyzing splicing regulatory motifs at *PTPRN2* locus, we extracted the ±10 bp sequence surrounding each variant and queried the sequence against the ATtRACT database (https://attract.cnic.es/seqsearch)^50^ to identify predicted RBP binding motifs overlapping the variant position.

### GWAS variants analysis

To predict splicing effects of disease-associated variants, we selected SNVs with nominal disease association (P < 1 × 10⁻⁵) from GWAS summary statistics. These included ALS GWAS data ^20^ comprising 29,612 cases and 122,656 controls of European and Asian ancestry; FTD GWAS data ^23^ comprising 2,154 cases and 4,308 controls of European ancestry; and the Pan-UK Biobank GWAS data ^22^ comprising six neurological or neuropsychiatric traits, including AD (839 cases and 410,833 controls, phecode-290.11), PD (1,660 cases and 403,249 controls, phecode-332), bipolar affective disorder (1,165 cases and 419,366 controls, icd10-F31), depression (527 cases and 369,930 controls, phecode-296.2), schizophrenia (680 cases and 419,906 controls, icd10-F20) and autism spectrum disease (186 cases and 135,760 controls, categorical-20544). Variants with Spliformer-V2 predicted Δscore > 0.1 were classified as splice-altering variants.

### Analysis of rare intronic variants in ALS WGS

To investigate rare intronic splice-altering variants in ALS, we analyzed 1,405 WGS data from four ALS cohorts: Chinese ALS (n = 307), Canadian ALS (n = 258), TargetALS (n = 164) and AnswerALS (n = 676) (**Table S3**). We used rare intronic variants (MAF < 0.0001) for Spliformer–V2 CSC/LMC model prediction.

### Tissue vulnerability analysis of rare splice-altering variants in ALS-associated genes

We used Spliformer-V2 to predict rare splice-altering variants across 18 tissues for SNVs located in 60 ALS-associated genes^12^ in 1,405 ALS WGS (**Table S3**). Variants with |Δscore| > 0.5 were classified as splice-altering. Enrichment of splice-altering variants in each tissue relative to other tissues was assessed using Fisher’s exact test.

### Evaluation of tissue specific splice-altering variants based on effect size

A reference mutation set was constructed following SpliceTransformer ^14^, using paired WGS and RNA-seq data from 181 frontal cortex, 157 LMC and 177 CSC samples. Splice-site usage was quantified using SpliSER v1.3. The final reference set comprised 855 SNVs with consistent directional splice-usage changes across all three tissues and |Δscore| > 0.1 in carriers versus noncarriers. Rare SNVs (MAF<0.0001) from ALS patients were then predicted using Spliformer-V2. For each tissue, SNVs with tissue Z-scores above the 95th percentile or below the 5th percentile of the reference Z-score distribution were considered to have stronger tissue-specific splicing effects.

### ClinVar data analysis

ClinVar variants were selected using the following criteria: the variant is an SNV; the variant mapped to only one gene; the variant was intronic or synonymous; and the variant was classified as benign or pathogenic. For each variant, Δscore was defined as the maximum donor or acceptor gain or loss score. We used Fisher’s exact test to compare the proportion of predicted splice-altering variants between pathogenic and benign groups

### Minigene assays, RT-PCR and amplicon length analysis

Minigene assays were used to validate candidate splice-altering variants in *PTPRN2* and *ERBB4*. We synthesized insert fragments containing exons and flanking intronic sequences (Shandong WZ Bioscience and Shanghai GENECHEM). Candidate variants were introduced using the KOD-Plus-Mutagenesis kit (TOYOBO, Japan), and all constructs were confirmed by Sanger sequencing.

SH-SY5Y cells were cultured in DMEM and supplemented with 10% fetal bovine serum (FBS, Gibco), Glutamax (Gibco), MEM non-essential amino acids solution (Gibco), and 1% penicillin/streptomycin (Gibco). Wild-type or mutant minigene plasmids were transfected using Lipo6000 transfection reagent (Beyotime, China). After 24–36 hours culture, total RNA was extracted and reverse-transcribed using the PrimeScript RT reagent kit (Takara Bio, Japan). RT-PCR was performed using 6-FAM-labeled primers (**Table S7)**, followed by amplicon-length analysis.

### Differential expression analysis of snRNA-seq data

Human motor cortex single-nucleus RNA-seq data were obtained from Synapse ID syn51105515 ^24^ (17 sporadic ALS cases and =16 controls). Seurat objects were processed for quality control, normalization, principal component analysis, integration and clustering. Final objects were merged using JoinLayers. Cell-type annotations were obtained from the provided metadata. Candidate gene expression was compared between ALS and control groups within each annotated cell type using the Wilcoxon rank-sum test.

### Differential expression and splicing analysis of RNA-seq data in TDP-43 knockdown neurons

Raw RNA-seq data from published TDP-43 knockdown neuron datasets^8, 25, 26^ were processed using fastp v0.23.2 for quality filtering and adapter trimming. Clean paired-end reads were aligned to hg38 using STAR v2.7.10b. Uniquely mapped reads were retained using samtools v1.21. Gene-level counts were generated using featureCounts from Rsubread v2.12.3 with GENCODE v48 annotation. Differential expression analysis was performed using DESeq2 v1.38.3 in R v4.2.2. P values were adjusted using the Benjamini-Hochberg method. For splicing analysis, we used LeafCutter to analyze the clean aligned paired reads for cryptic splicing events detection.

### Differential expression analysis of RNA-seq data in rat TDP-43 mutant neurons

RNA-seq data from rat hippocampal neurons expressing TDP-43 mutants A90V, M337V or Q331K, or GFP control, were obtained from our previous study^12^. Reads were processed as previously described and aligned to the rat reference genome mRatBN7.2/rn7. Gene-level read counts were generated using featureCounts v2.0.1. Differential expression analysis was performed using DESeq2 v1.44.0, comparing each TDP-43 mutant condition with GFP control.

### Association between PTPRN2 expression and ALS clinical phenotypes

We estimated *PTPRN2* mRNA expression from RNA-seq data of TargetALS spinal cord and motor cortex tissues. Multivariate linear regression was used to test associations between *PTPRN2* expression, defined as log₂(TPM + 1), and disease duration or age at symptom onset, adjusting for sex.

### *PTPRN2* knockdown and RT-qPCR in SH-SY5Y cells

Cells were transfected with control shRNA, shPTPRN2 for 48 hours (**Table S8**). Total RNA was extracted and reverse-transcribed using the PrimeScript RT reagent kit (Takara Bio, Japan). qPCR was performed on a QuantStudio 7 Flex real-time PCR system using TB Green Premix Ex Taq (Takara Bio, Shiga, Japan) and primers listed in **Table S9**. We used unpaired two-tailed Student’s t-test to measure mRNA expression alteration. Statistical analysis and visualization were performed in GraphPad Prism v9.5.

### Immunofluorescence analysis of TDP-43 subcellular localization

SH-SY5Y cells were seeded in confocal dishes and transfected with control shRNA, shPTPRN2, or shPTPRN2 together with pCMV-PTPRN2(human)-P2A-mCherry-Neo plasmid (Miaoling Biology, China) for rescue experiments (**Table S8**). After 48 hours culture, cells were fixed in 4% paraformaldehyde, permeabilized with 0.5% Triton X-100 and blocked with 1% BSA. Cells were incubated overnight at 4 °C with anti-TDP-43 antibody (Proteintech, USA, 1:2000), followed by AF647-conjugated goat anti-rabbit or AF555-conjugated donkey anti-rabbit IgG (H+L) secondary antibody (Beyotime, China, 1:500) for 1 h at room temperature in the dark. Nuclei were counterstained with DAPI. Images were acquired using a Nikon TI2-E+A1 inverted confocal microscope. Five random fields of view were analyzed per condition. The proportion of cells with cytoplasmic TDP-43 mislocalization was compared using one-way ANOVA followed by Tukey’s multiple-comparison test.

### siRNA transfection in iPSC-induced neurons (iNs)

The iPSC line containing a stable integration of TetO-Ngn2 was a generous gift from Dr. Martin Kampmann at UCSF^51^. Cells were maintained in mTeSR1 medium (Stem Cell Technologies, Cat #85850) on Matrigel-coated plates (BD, Cat #8774552) at 37°C in 5% CO₂. When cultures reached approximately 80% confluence, the medium was switched to induction medium (IM), consisting of DMEM/F12 (Corning, Cat #10-090-CV), GlutaMAX (Gibco, Cat #35050061), N2 (Gibco, Cat #17502048), NEAA (Gibco, Cat #11140035), and penicillin/streptomycin (Corning, Cat #MT30001CI) supplemented with ROCK inhibitor (Cat #S6390) and 1 μg/mL doxycycline (Dox, Cayman, Cat #14422) to induce NGN2 expression. On Day 2, the medium was completely replaced with fresh IM containing ROCK inhibitor, 1 μg/mL Dox, 10 ng/mL BDNF (R&Dsystem, Cat #11166-BD), and 10 ng/mL NT3. On Day 3, cells were dissociated and replated onto Matrigel-coated plates in the same medium supplemented with 40 μM BrdU (Cayman Cat #15580). On Day 4, the medium was replaced with N3 neuronal medium containing 1 μg/mL Dox, 10 ng/mL BDNF, 10 ng/mL NT3. On Day 5, cells were transfected with siRNAs (**Table S8**) at a final concentration of 30 nM using Lipofectamine RNAiMAX transfection reagent (ThermoFisher, Cat #13778150). The medium was completely replaced 24 h after transfection. Cells were harvested 72 h after transfection for qRT-PCR (**Table S10**) and western blot analyses (anti-TDP-43 antibody, 1:1000, Proteintech, Cat #10782-2-AP; anti-PTPRN2 antibody, 1:1000, MilliporeSigma, Cat #HPA006900-100UL).

### Molecular cloning and viral production

Virus vector plasmids containing iMN factors (Ngn2, Lhx3, Isl1, NeuroD1, Ascl1, Brn2 and Myt1l) and Hb9:RFP were generated as previously described 8. SYN1:GFP and SYN1:PTPRN2:GFP lentiviral vector plasmids were purchased from VectorBuilder (ID: VB250911-1748ynn, VB250804-1477sfw). Viruses were generated as follows. iMNs factor or target gene viral vectors, together with viral packaging plasmids (pPAX2 and VSVG for lentivirus; PLK and pHDM for retrovirus), were transfected to HEK293T cells cultured in a 10-cm dish at 80-90% confluence using polyethylenimine (PEI, Kyfora Bio, Cat #23966-1). The culture medium was replaced 24 h after transfection. Virus-containing supernatant was collected at 48 h and 72 h after transfection. The collected supernatant was passed through 4.5-um filters, incubated with lenti-X (Takara, Cat #631232) overnight at 4°C, and then centrifuged at 1,500 g for 45 min at 4°C. Virus pellets were resuspended in DMEM containing 10% FBS (200 ul per 10-cm dish) and stored at −80°C until use.

### Conversion of fibroblast-like cells into induced motor neurons (iMNs)

Conversion of fibroblast-like cells into iMNs was performed as previously described ^52^ with minor modifications. Fibroblasts derived from patient iPSCs were seeded onto 96-well plates coated with 0.1% gelatin (10 min at RT) followed by laminin (ThermoFisher, Cat #23017015, overnight at 4°C) at a density of 3000-4000 cells per well. On Day 0, fibroblasts were infected with seven iMN factor retroviruses in the presence of 8 μg/mL polybrene (Sigma, Cat #H9268-5G). On Day 2, Hb9:RFP lentivirus was added to the cultures. On Day 4, cultures were infected with either SYN1:PTPRN2:GFP or SYN1:GFP lentivirus. Complete medium changes were performed 24 h after each viral infection. On Day 6, primary mouse cortical glial cells isolated from P2–P3 wild-type pups were added to the cultures in glial medium consisting of DMEM/F12, GlutaMAX, 10% FBS (GenClone, Cat #25-514) and penicillin/streptomycin. On Day 7, the glial medium was replaced with N3 neuronal medium containing DMEM/F12, 5% FBS, GlutaMAX, N2 and B27 (Gibco, Cat #17502048 and #12587010), 7.5 μM RepSox (Selleckchem, Cat #S7223), and 10 ng/mL each of FGF2 (ThermoFisher, 100-18B-100UG), BDNF, GDNF, and CNTF (R&Dsystems, Cat #11166-BD-050, #212-GD-050/CF, and #257-NT-050/CF). The cultures were maintained in N3 medium with medium changes every other day.

## Survival assay

Hb9:RFP-positive iMNs typically formed between days 13 and 16. Survival assays began on Day 17 by withdrawing neurotrophic factors (RepSox, FGF2, BDNF, GDNF, and CNTF) from the N3 medium to exacerbate the survival differences between healthy control lines (two lines) and ALS/FTD patient lines (three lines) (**Table S11**). Individual iMNs were longitudinally tracked every other day for 13 days, and all images were analyzed using ImageJ. Only GFP+/RFP+ iMNs were included in the survival analysis. Cells were manually scored as dead when their somas fragmented into two or three detached parts, as indicated by the RFP signal. Survival analyses were performed using the Kaplan–Meier method. Survival curves were compared using the log-rank (Mantel–Cox) test, and hazard ratios were calculated using the Mantel–Haenszel method. For hazard ratio analyses, the hazard rate of healthy control iMNs expressing GFP was normalized to 1, and the hazard rates of all other conditions were divided by the control hazard rate to calculate hazard ratios. Statistical significance among hazard ratios was determined by one-way ANOVA.

## Data availability

Raw WGS and RNA-seq data of CSC and LMC tissues were obtained from the TargetALS Human Postmortem Tissue Core, the New York Genome Center for Genomics of Neurodegenerative Disease, the ALS Association and the Tow Foundation; GTEx v10 raw WGS and RNA-seq data of 16 human tissues in the model development were obtained from dbGaP accession phs000424.v10.p2(https://www.ncbi.nlm.nih.gov/projects/gap/cgi-bin/study.cgi?study_id=phs000424.v10.p2), under the project approval (#30134) of ‘Development of a novel deep learning tool to predict pre-mRNA splicing’. Pathogenic and benign variants were obtained from ClinVar (https://ftp.ncbi.nlm.nih.gov/pub/clinvar/vcf_GRCh38/clinvar.vcf.gz). Human sQTL data across tissues were obtained from GTEx v10 single-tissue cis-QTL resource (https://storage.googleapis.com/adult-gtex/bulk-qtl/v10/single-tissue-cis-qtl/GTEx_Analysis_v10_sQTL.tar). The Ascot data was derived from http://ascot.cs.jhu.edu.

ALS GWAS summary results were obtained from https://projectmine.com/research/^20^, FTD GWAS summary results were obtained from a previous study ^23^, GWAS summary results from AD, PD, BPD, depression, autism and schizophrenia were obtained from https://pan.ukbb.broadinstitute.org/phenotypes. Raw WGS data from the Canadian ALS patients is part of the MinE project and available upon reasonable data request (https://www.projectmine.com/research/data-sharing/). Raw ALS WGS data from western populations were obtained from the AnswerALS portal, and the TargetALS. Rare variants data from WGS of Chinese ALS patients (GVM accession: GVM000544) were deposited in the Genome Sequence Archive for human and Genome Variation Map (GVM) in the National Genomics Data Center, China National Center for Bioinformation/Beijing Institute of Genomics, Chinese Academy of Sciences (https://ngdc.cncb.ac.cn/). ALS RNA-seq data from spinal cord and motor cortex tissues were obtained from the TargetALS.

## Code availability

Spliformer-V2 code is available at GitHub: https://github.com/TJ-zhanglab/Spliformer-V2, which includes codes from model inference. The model train, test and benchmarking code will be deposited at Zenodo. The LMC and CSC model weights will be deposited in Hugging Face: https://huggingface.co/tjzhanglab. Spliformer-V2 model weights of other tissues are not publicly available according to the GTEx policy.

## Author contributions

M.Z. conceived and designed the study; X.L.T, and H.L.L contributed to data preparation, model development, and model evaluations; M.S. contributed on the work of iPSC neurons; X.Y.M worked on TDP-43 localization experiments; J.Y.G worked on snRNA-seq analysis; X.L.T, H.L.L and Y.F.S analyzed WGS and RNA-seq data and the public databases; X.L.T and X.Y.M worked on minigene assays; E.R, L.Z and A.A contributed to Canadian ALS samples and WGS data; Y.C and Y.D contributed to Chinese ALS samples and clinical data; Q.H.W contributed to cell-based immunofluorescence work; Y.L.C contributed to RNA-seq data of rat neurons; A.D.G and Y.Z provided RNA-seq data of iNeurons; X.L.T, H.L.L, X.Y.M and M.S worked on drafting graphs and tables in the manuscript; M.Z, J.K.I, Y.C provided data, laboratory, computational and clinical resources to the project; M.Z and J.K.I. supervised the project; M.Z, X.L.T, M.S and J.K.I. worked on the manuscript and revisions.

## Acknowledgements

We would like to thank the ‘GTEx project’, ‘Target ALS Human Postmortem Tissue Core’, ‘New York Genome Center for Genomics of Neurodegenerative Disease’, and the ‘Amyotrophic Lateral Sclerosis Association and the Tow Foundation’, the ‘MinE project’, the ‘UK biobank (project number: 256056)’ for providing multi-omics dataset; and ALS patients contributing samples relevant to this project.

## Reference

1. Rogalska, M.E., Vivori, C. & Valcárcel, J. Regulation of pre-mRNA splicing: roles in physiology and disease, and therapeutic prospects. Nat Rev Genet 24, 251–269 (2023).

2. Angarola, B.L. & Anczuków, O. Splicing alterations in healthy aging and disease. Wiley Interdiscip Rev RNA 12, e1643 (2021).

3. Maurano, M.T. et al. Systematic localization of common disease-associated variation in regulatory DNA. Science 337, 1190–1195 (2012).

4. Consortium, G.T. The GTEx Consortium atlas of genetic regulatory effects across human tissues. Science 369, 1318–1330 (2020).

5. Shukla, S. & Oberdoerffer, S. Co-transcriptional regulation of alternative pre-mRNA splicing. Biochim Biophys Acta 1819, 673–683 (2012).

6. Fu, X.D. & Ares, M., Jr. Context-dependent control of alternative splicing by RNA-binding proteins. Nat Rev Genet 15, 689–701 (2014).

7. Ling, J.P., Pletnikova, O., Troncoso, J.C. & Wong, P.C. TDP-43 repression of nonconserved cryptic exons is compromised in ALS-FTD. Science 349, 650–655 (2015).

8. Klim, J.R. et al. ALS-implicated protein TDP-43 sustains levels of STMN2, a mediator of motor neuron growth and repair. Nat Neurosci 22, 167–179 (2019).

9. Brown, A.L. et al. TDP-43 loss and ALS-risk SNPs drive mis-splicing and depletion of UNC13A. Nature 603, 131–137 (2022).

10. Ma, X.R. et al. TDP-43 represses cryptic exon inclusion in the FTD-ALS gene UNC13A. Nature 603, 124–130 (2022).

11. Jaganathan, K. et al. Predicting Splicing from Primary Sequence with Deep Learning. Cell 176, 535–548 e524 (2019).

12. Tang, X. et al. Deep learning analyses of splicing variants identify the link of PCP4 with amyotrophic lateral sclerosis. Brain, awaf025 (2025).

13. Zeng, T. & Li, Y.I. Predicting RNA splicing from DNA sequence using Pangolin. Genome Biol 23, 103 (2022).

14. You, N. et al. SpliceTransformer predicts tissue-specific splicing linked to human diseases. Nat Commun 15, 9129 (2024).

15. Dalla-Torre, H. et al. Nucleotide Transformer: building and evaluating robust foundation models for human genomics. Nat Methods 22, 287–297 (2025).

16. de Almeida, B.P. et al. SegmentNT: annotating the genome at single-nucleotide resolution with DNA foundation models. bioRxiv, 2024.2003. 2014.584712 (2024).

17. Wagner, N. et al. Aberrant splicing prediction across human tissues. Nat Genet 55, 861–870 (2023).

18. Moakley, D.F. et al. Reverse engineering neuron-type-specific and type-orthogonal splicing-regulatory networks using diverse cellular transcriptomes. Cell Rep 44, 115898 (2025).

19. Dasgupta, T. & Ladd, A.N. The importance of CELF control: molecular and biological roles of the CUG-BP, Elav-like family of RNA-binding proteins. Wiley Interdiscip Rev RNA 3, 104–121 (2012).

20. van Rheenen, W. et al. Common and rare variant association analyses in amyotrophic lateral sclerosis identify 15 risk loci with distinct genetic architectures and neuron-specific biology. Nat Genet 53, 1636–1648 (2021).

21. Guo, Y. et al. SMR-Portal: an online platform for integrative analysis of GWAS and xQTL data to identify complex trait genes. Nat Methods 22, 220–222 (2025).

22. Karczewski, K.J. et al. Pan-UK Biobank GWAS improves discovery, analysis of genetic architecture, and resolution into ancestry-enriched effects. medRxiv, 2024.2003.2013.24303864 (2024).

23. Ferrari, R. et al. Frontotemporal dementia and its subtypes: a genome-wide association study. Lancet Neurol 13, 686–699 (2014).

24. Pineda, S.S. et al. Single-cell dissection of the human motor and prefrontal cortices in ALS and FTLD. Cell 187, 1971–1989 e1916 (2024).

25. Brown, A.L., et al. Sensitivity to TDP-43 loss and degradation resistance determine cryptic exon biomarker potential. *bioRxiv* (2025).

26. Zeng, Y., et al. Nonsense-mediated decay masks cryptic splicing events caused by TDP-43 loss. *bioRxiv* (2025).

27. Vu, L. et al. Proteomics and mathematical modeling of longitudinal CSF differentiates fast versus slow ALS progression. Annals of clinical and translational neurology 10, 2025–2042 (2023).

28. van der Ende, E.L. et al. Novel CSF biomarkers in genetic frontotemporal dementia identified by proteomics. Annals of clinical and translational neurology 6, 698–707 (2019).

29. Mravinacová, S. et al. Addressing inter individual variability in CSF levels of brain derived proteins across neurodegenerative diseases. Scientific reports 15, 668 (2025).

30. Seddighi, S. et al. Mis-spliced transcripts generate de novo proteins in TDP-43–related ALS/FTD. Science translational medicine 16, eadg7162 (2024).

31. Mathur, M. et al. Programmable mutually exclusive alternative splicing for generating RNA and protein diversity. Nat Commun 10, 2673 (2019).

32. Olivieri, J.E. et al. RNA splicing programs define tissue compartments and cell types at single-cell resolution. eLife 10, e70692 (2021).

33. Cheng, S.J. et al. Accurately annotate compound effects of genetic variants using a context-sensitive framework. Nucleic Acids Res 45, e82 (2017).

34. Yeo, G., Holste, D., Kreiman, G. & Burge, C.B. Variation in alternative splicing across human tissues. Genome Biol 5, R74 (2004).

35. Nikom, D. & Zheng, S. Alternative splicing in neurodegenerative disease and the promise of RNA therapies. Nat Rev Neurosci 24, 457–473 (2023).

36. Guo, C. et al. Cryptic splicing in synaptic and membrane excitability genes links TDP-43 loss to neuronal dysfunction. Sci Transl Med 18, eaeb8517 (2026).

37. Joseph, B.J. et al. TDP-43-dependent mis-splicing of KCNQ2 triggers intrinsic neuronal hyperexcitability in ALS/FTD. Nat Neurosci 28, 2476–2492 (2025).

38. Xiao, S. et al. RNA targets of TDP-43 identified by UV-CLIP are deregulated in ALS. Molecular and Cellular Neuroscience 47, 167–180 (2011).

39. Yang, Y.-C.T. et al. CLIPdb: a CLIP-seq database for protein-RNA interactions. BMC genomics 16, 51 (2015).

40. Baughn, M.W. et al. Mechanism of STMN2 cryptic splice-polyadenylation and its correction for TDP-43 proteinopathies. Science 379, 1140–1149 (2023).

41. Delaneau, O., Zagury, J.F., Robinson, M.R., Marchini, J.L. & Dermitzakis, E.T. Accurate, scalable and integrative haplotype estimation. Nat Commun 10, 5436 (2019).

42. Browning, B.L., Tian, X., Zhou, Y. & Browning, S.R. Fast two-stage phasing of large-scale sequence data. Am J Hum Genet 108, 1880–1890 (2021).

43. Joglekar, A. et al. Single-cell long-read sequencing-based mapping reveals specialized splicing patterns in developing and adult mouse and human brain. Nat Neurosci 27, 1051–1063 (2024).

44. Zhou, Z. et al. HELIX: a scalable model for predicting context-dependent regulation of RNA splicing and isoform usage. Nat Comput Sci 6, 464–477 (2026).

45. Celaj, A. et al. An RNA foundation model enables discovery of disease mechanisms and candidate therapeutics. BioRxiv, 2023.2009. 2020.558508 (2023).

46. Mertes, C. et al. Detection of aberrant splicing events in RNA-seq data using FRASER. Nat Commun 12, 529 (2021).

47. Ling, J.P. et al. ASCOT identifies key regulators of neuronal subtype-specific splicing. Nat Commun 11, 137 (2020).

48. Avsec, Ž., et al. Advancing regulatory variant effect prediction with AlphaGenome. Nature 649, 1206–1218 (2026).

49. Grant, C.E., Bailey, T.L. & Noble, W.S. FIMO: scanning for occurrences of a given motif. Bioinformatics 27, 1017–1018 (2011).

50. Giudice, G., Sanchez-Cabo, F., Torroja, C. & Lara-Pezzi, E. ATtRACT-a database of RNA-binding proteins and associated motifs. Database (Oxford) 2016 (2016).

51. Tian, R. et al. CRISPR interference-based platform for multimodal genetic screens in human iPSC-derived neurons. Neuron 104, 239–255. e212 (2019).

52. Linares, G.R. et al. SYF2 suppression mitigates neurodegeneration in models of diverse forms of ALS. Cell Stem Cell 30, 171–187. e114 (2023).

