## Supplementary material for "Spliformer-V2 enables multi-tissue prediction and interpretation of splice-altering genetic variants"

### Supplementary materials

#### Supplementary Figures

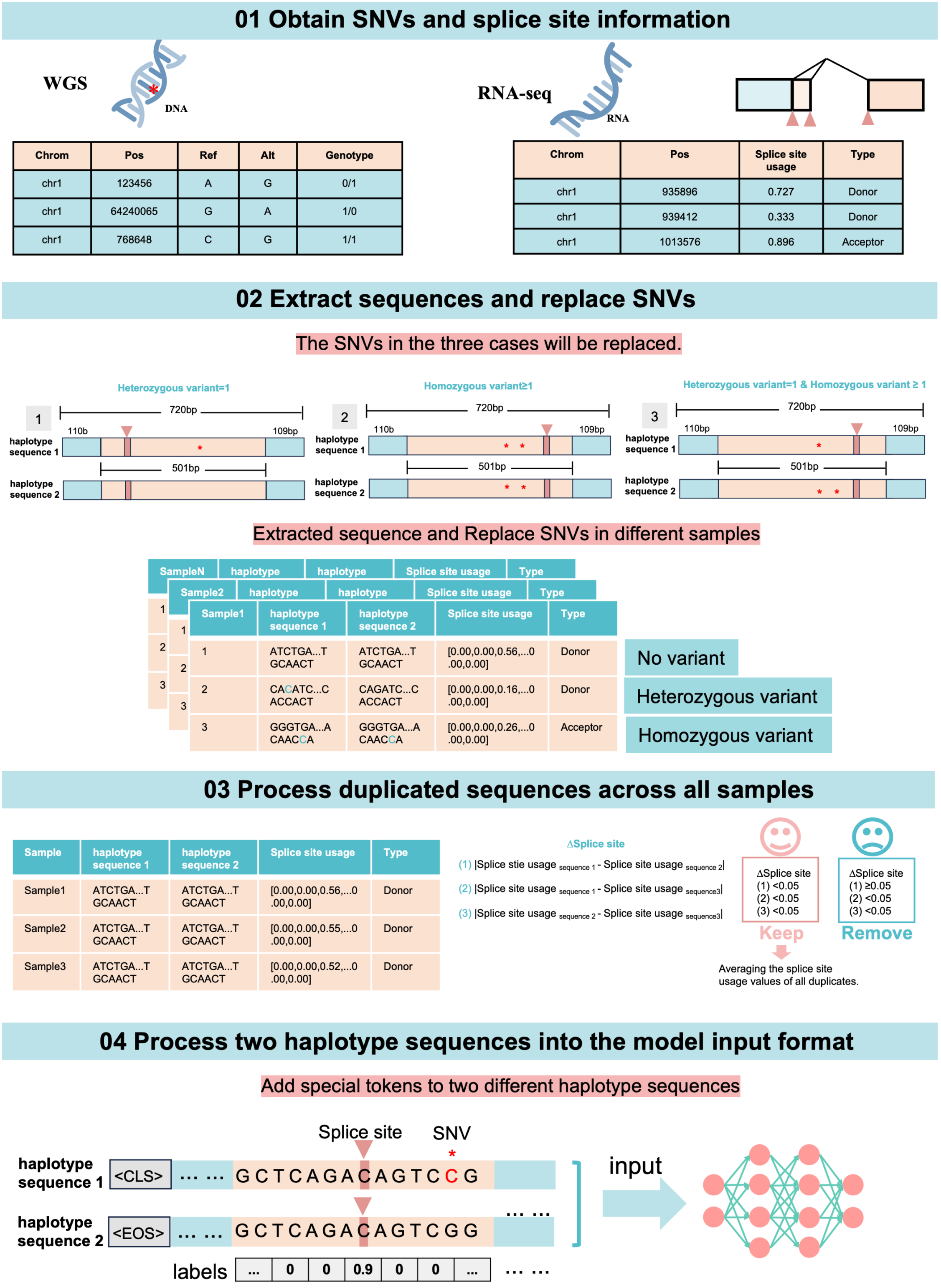

**Fig. S1 Workflow for dataset preparation.** The procedure consists of four main steps. First, SNV information was extracted from TargetALS/GTEx VCF files. RNA-seq data from 18 tissues across 20 individuals were aligned to the human reference genome (hg38), and splice site usage was quantified for each tissue. Second, a 720 bp reference sequence with a splice site at the center or at the other position was extracted, and SNVs were replaced according to genotype. Only SNVs meeting one of the following criteria were replaced: (i) a single heterozygous variant within ±360 bp; (ii) one or more homozygous variants within ±360 bp; (iii) both variant types within ±360 bp, with only one heterozygous and any number of homozygous variants allowed. Third, all sequences from the 20 individuals were merged. For each group of duplicate sequences, pairwise differences in splice site usage were calculated. Only groups with all differences < 0.05 were retained, and the final label was the average usage across duplicates. Finally, the two haplotype sequences were concatenated into a single input sequence (diploid sequence) by adding special tokens: <CLS> at the beginning of the first haplotype sequence and <EOS> at the beginning of the second haplotype sequence.

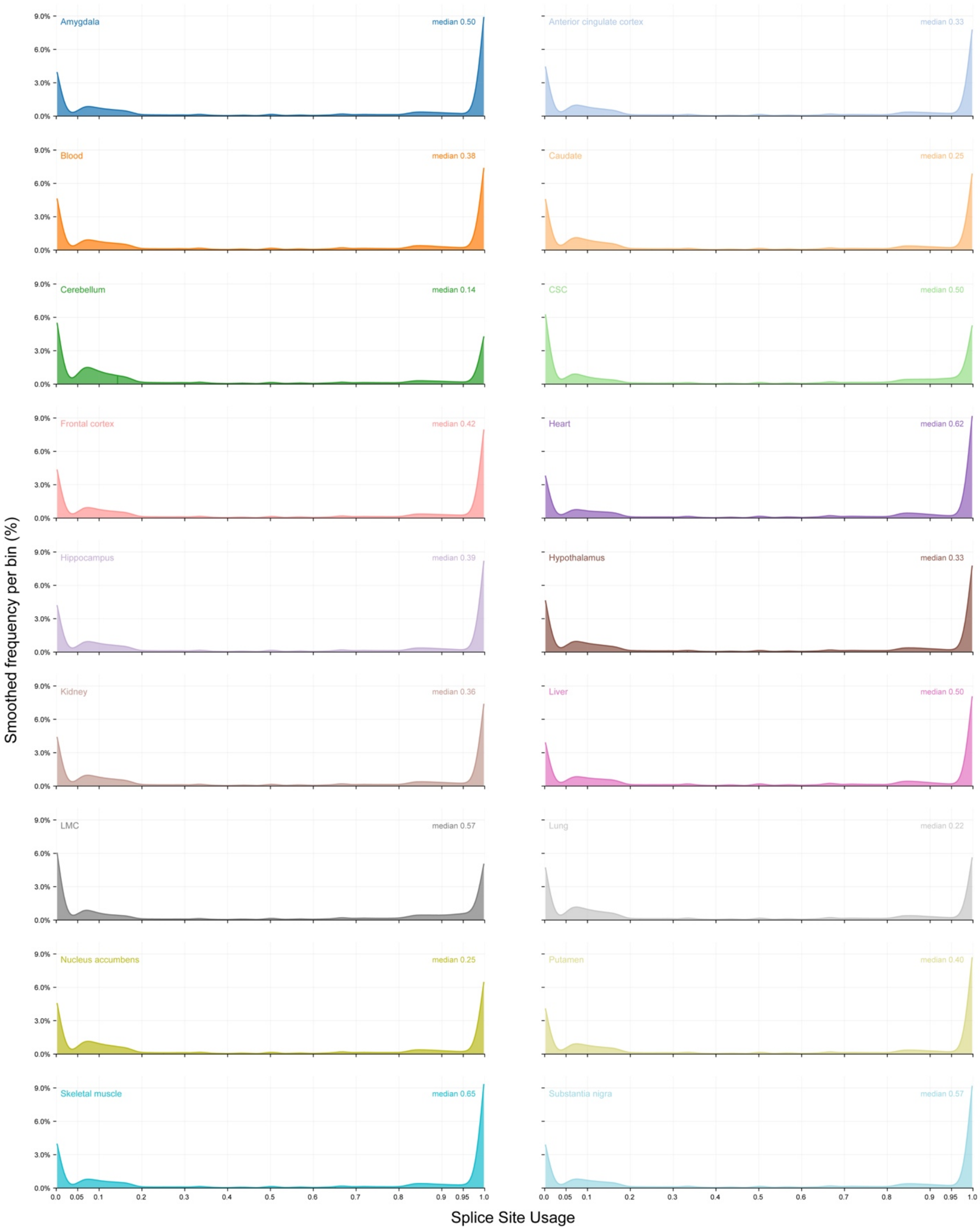


**Fig S2. Distribution of splice-site usage across tissues.** The x-axis represents splice-site usage, and the y-axis represents the smoothed frequency of records within each usage bin (bin width= 0.004). The median splice site usage values were shown for each tissue. Sequences were extracted using the center-window approach.

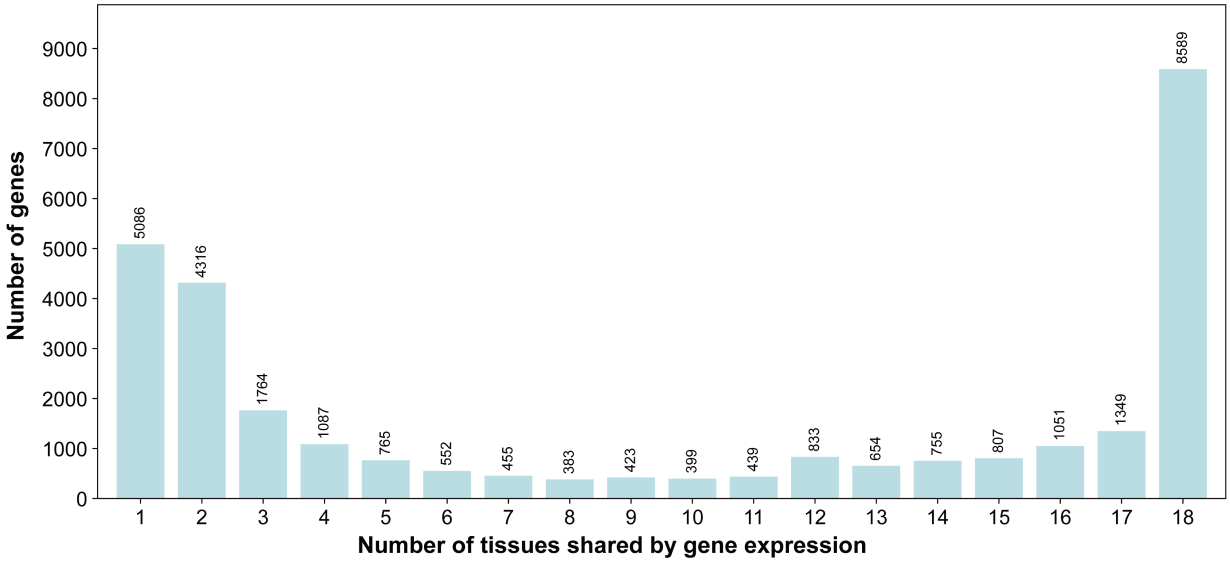


**Fig S3. Histogram showing the distribution of number of genes shared across different number of tissues based on gene expression.** It includes protein-coding genes, lncRNA, miRNA, and snRNA. A gene was considered expressed in a tissue when its median TPM across all samples from that tissue was ⩾ 1. Genes expressed in 1–18 tissues are shown, with numbers above the bars indicating the number of genes expression shared across tissues. Gene-level counts were generated using featureCounts v2.1.1 and normalized to transcripts per million (TPM) using featureCounts-reported gene lengths in the multi-tissue dataset.

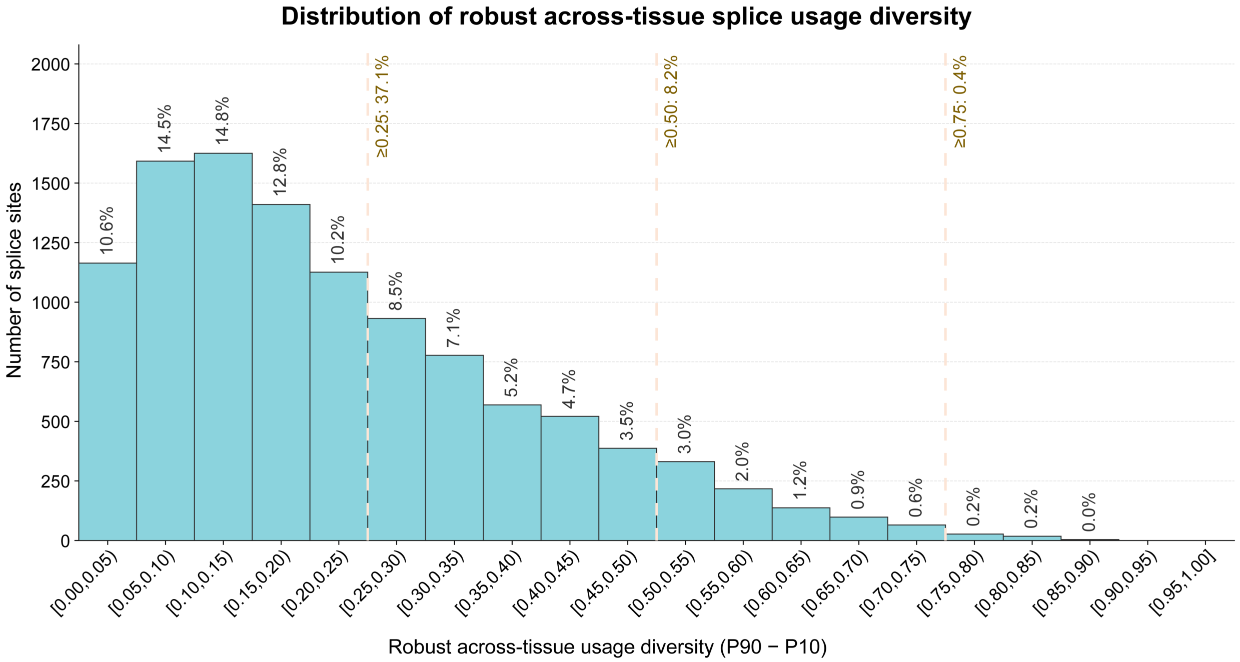


**Fig S4. Histogram showing the distribution of robust across-tissue splicing usage diversity.** A total of 11,000 splice sites detected in at least 14 of 18 tissues were included. For each splice site, all available usage values within each tissue were first averaged, and robust across-tissue usage diversity was then defined as the difference between the 90th and 10th percentiles of the tissue-level mean usage values (P90−P10). Vertical dashed lines mark three reference thresholds together with the percentages of splice sites at or above each threshold: 0.25 (37.1%), 0.50 (8.2%), and 0.75 (0.4%). Linked to **Fig.1F**

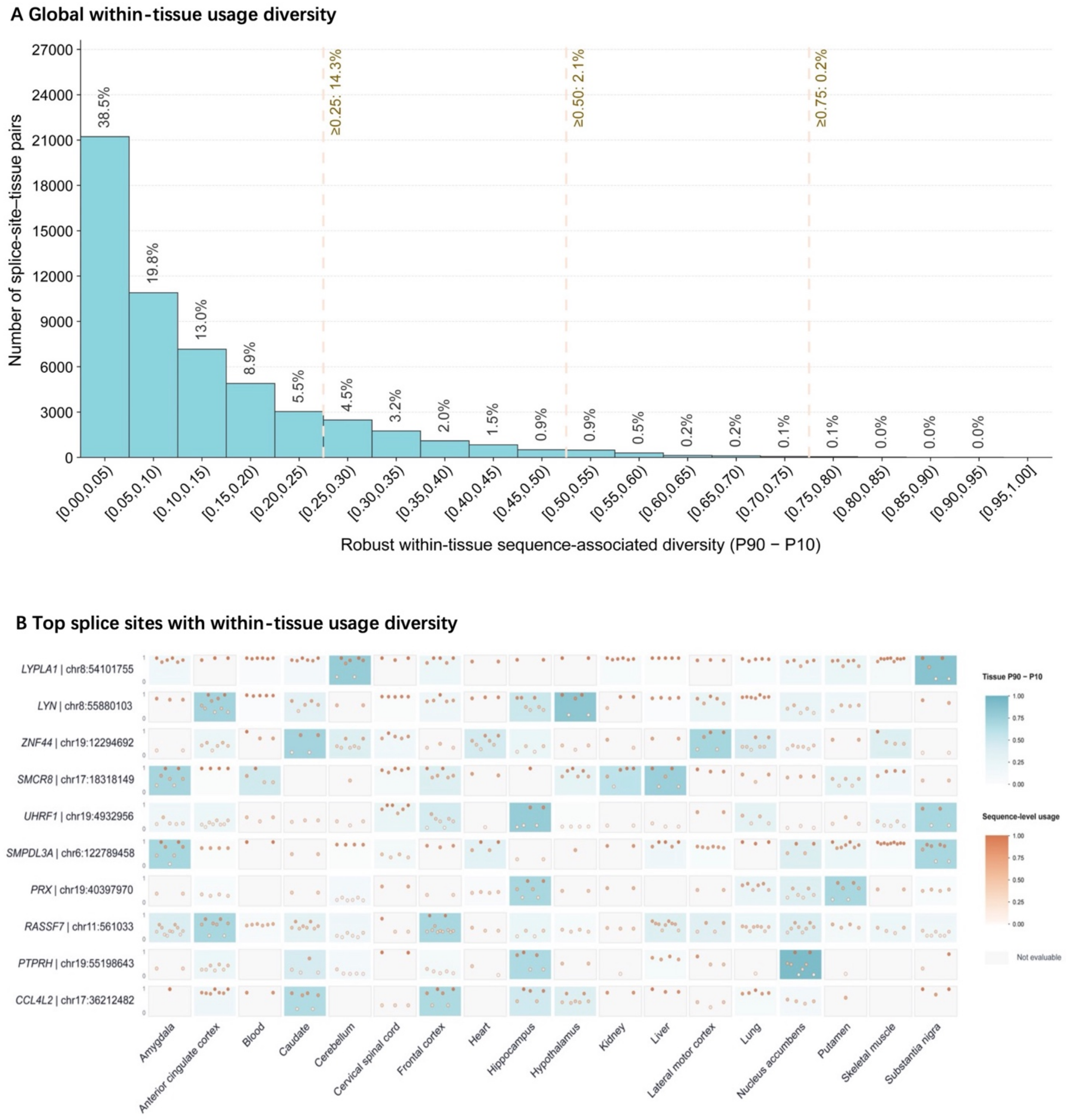


**Fig. S5. Within-tissue sequence-associated diversity of splicing usage. (A)** Histogram showing the global distribution of robust within-tissue sequence-associated usage diversity across 55,093 evaluable splice-site–tissue pairs. The analysis included splice sites with usage information in at least 14 of 18 tissues. For each splice site in each tissue, at least four sequence-level usage values were required, and robust diversity was calculated as the difference between the 90th and 10th percentiles of these values (P90−P10). Vertical dashed lines mark three reference thresholds together with the percentages of pairs at or above each threshold: 0.25 (14.3%), 0.50 (2.1%), and 0.75 (0.2%). **(B)** Heatmap showing ten splice sites with recurrently high within-tissue sequence-associated diversity. Splice sites were required to have at least six evaluable tissues and were ranked by the median of their three largest tissue-specific P90−P10 values. Rows represent splice sites and columns represent tissues. Blue tile intensity represents P90−P10 diversity calculated from the sequence-level usage observations for the indicated splice site and tissue. Near-white tiles denote non-evaluable pairs with fewer than four available usage values. Peach points show all available sequence-level usage values; both their vertical positions and colors correspond to usage on a 0–1 scale, with 0 at the bottom and 1 at the top of each tile. At right, the blue color key represents tile-level P90−P10 diversity, and sequence-level usage diversity that may contain one or multiple variants.

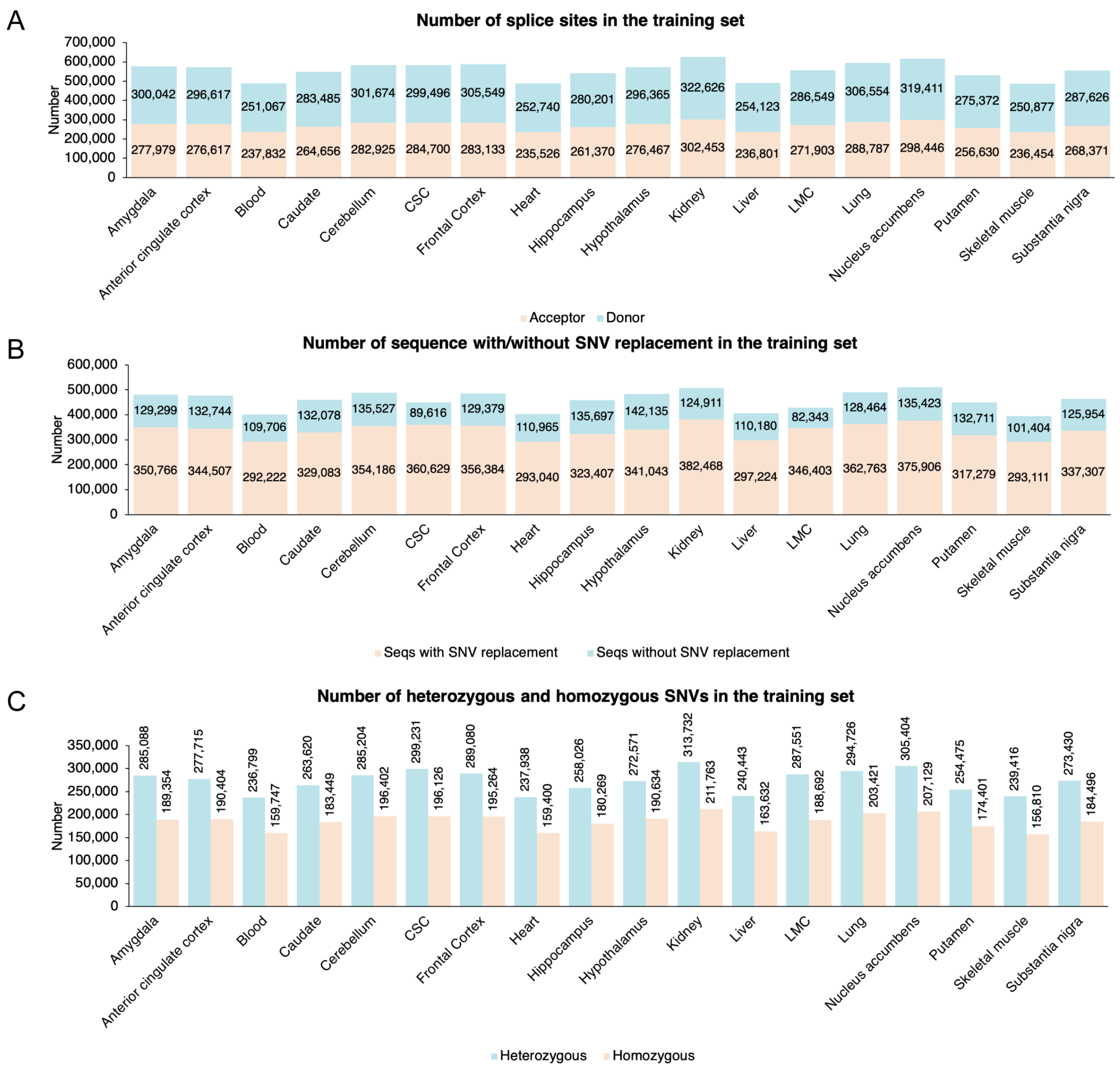


**Fig. S6 Details of the training set. (A)** Number of donor and acceptor sites in the training set **(B)** Number of sequences with/without SNV replacement in the training set. **(C)** Number of heterozygous and homozygous SNVs in the training set across 18 tissues.

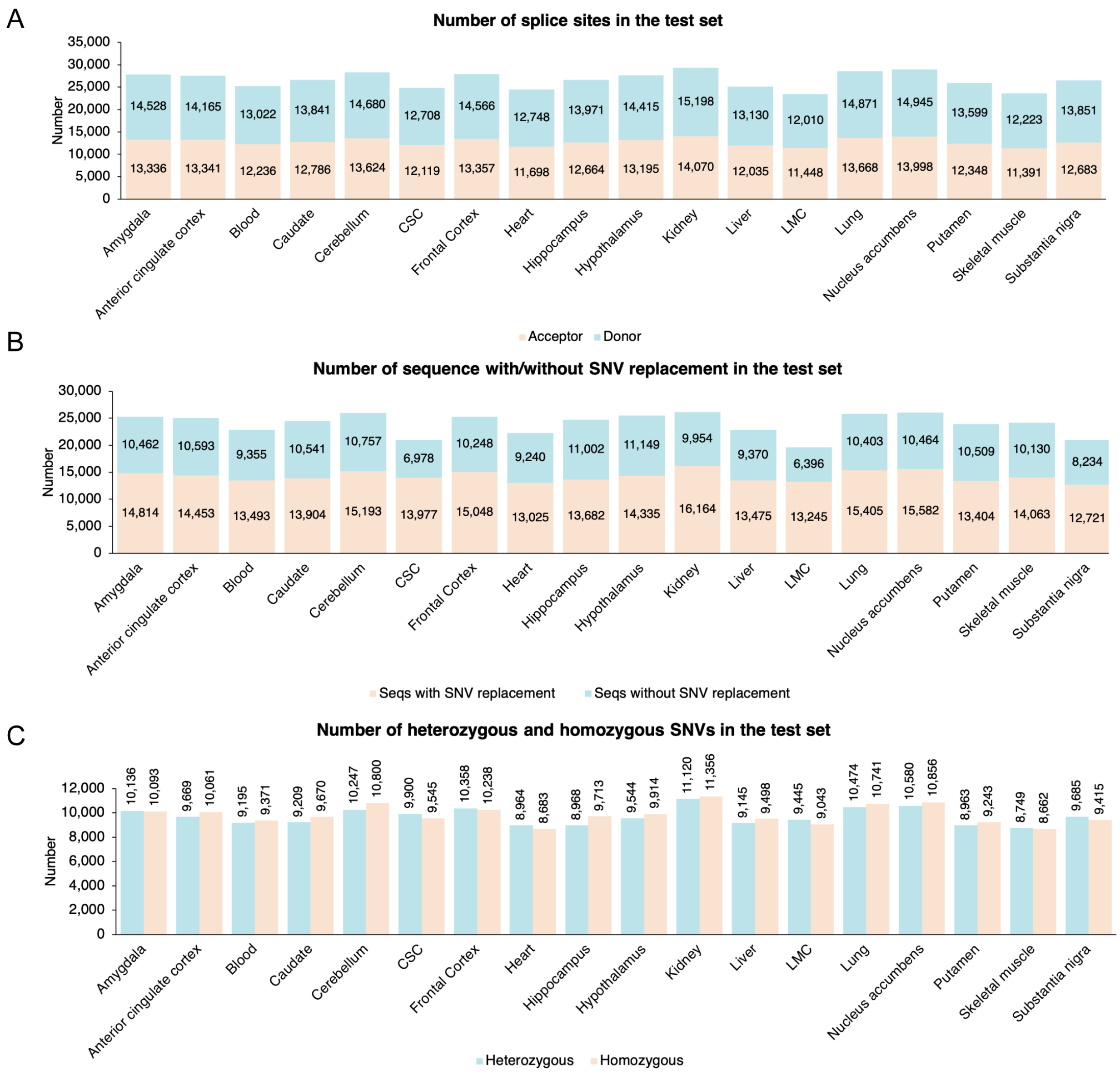


**Fig. S7 Details of the test set. (A)** Number of donor and acceptor sites in the test set **(B)** Number of sequences with/without SNV replacement in the test set. **(C)** Number of heterozygous and homozygous SNVs in the test set across 18 tissues.

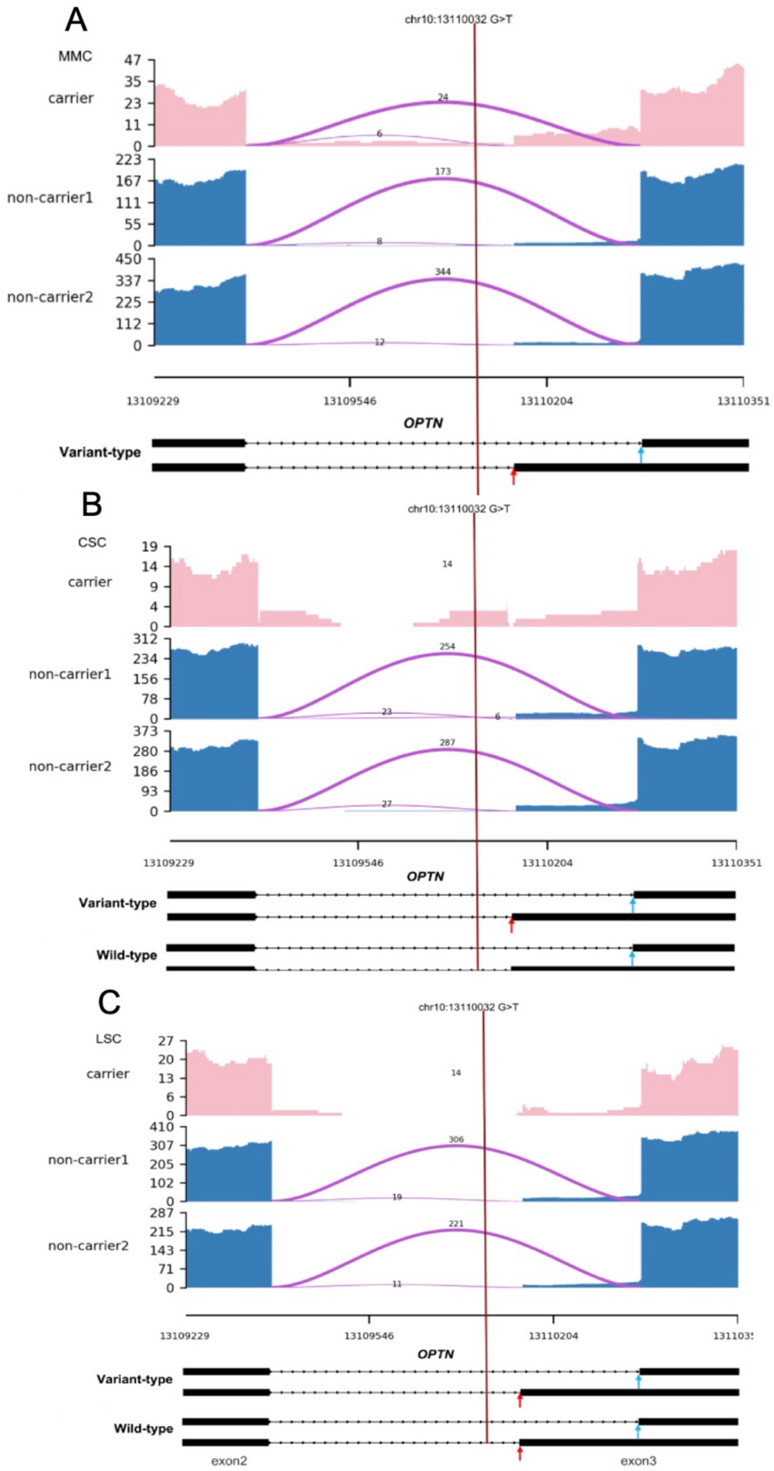


**Fig. S8** Visualization of the splicing effect of a variant located in *OPTN* (chr10:13110032 G>T, hg38) in MMC (**A**), CSC (**B**) and LSC (**C**). In all three tissues, this variant was not identified as an outlier by FRASER.

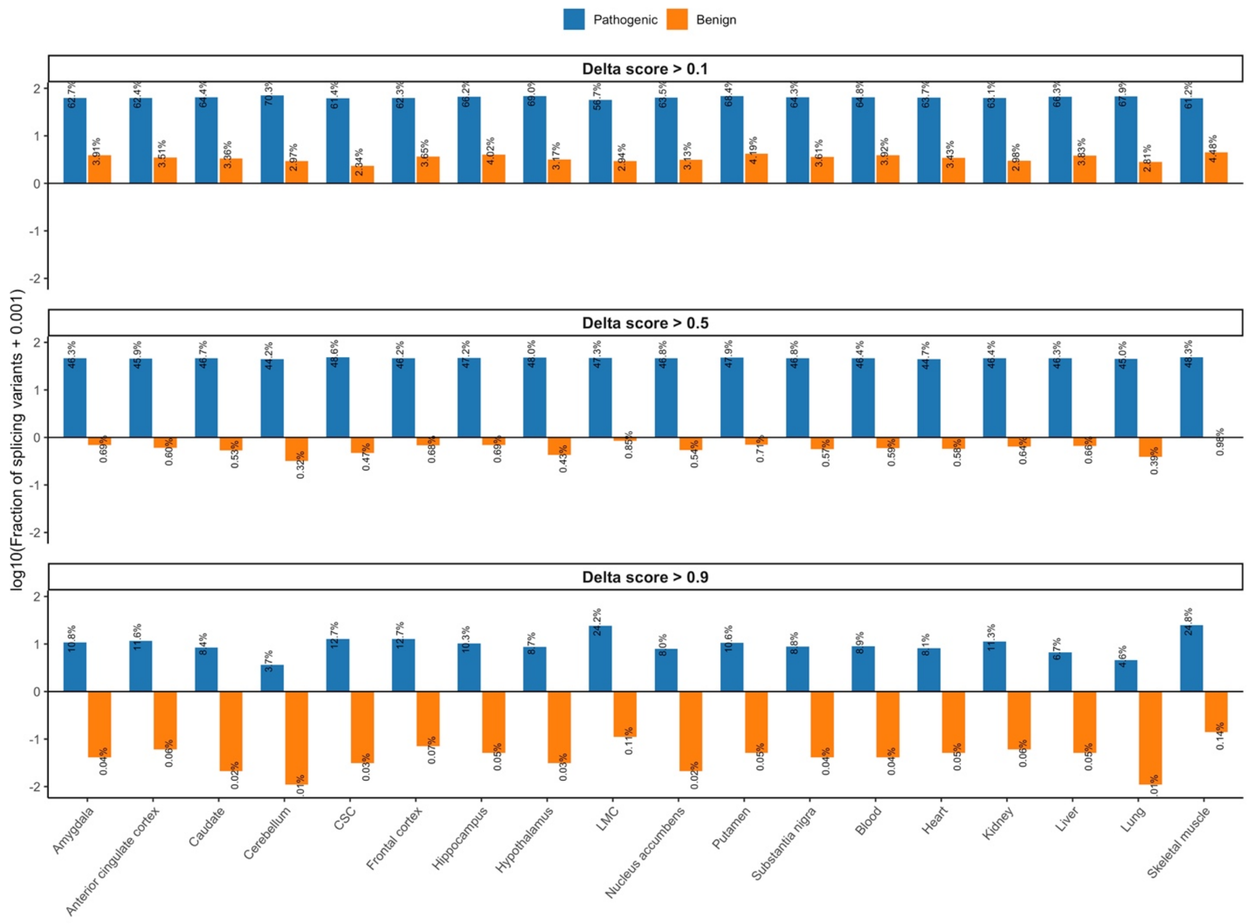


**Fig. S9 Spliformer-V2 analysis of ClinVar dataset reveals a link between splicing variants and disease pathogenicity.** The bar plots showing log10 transformation of the fraction of splicing variants in pathogenic or benign variants at different delta score cutoffs across 18 tissues. The actual fraction of splicing variants (percentage) was labeled for each bar.

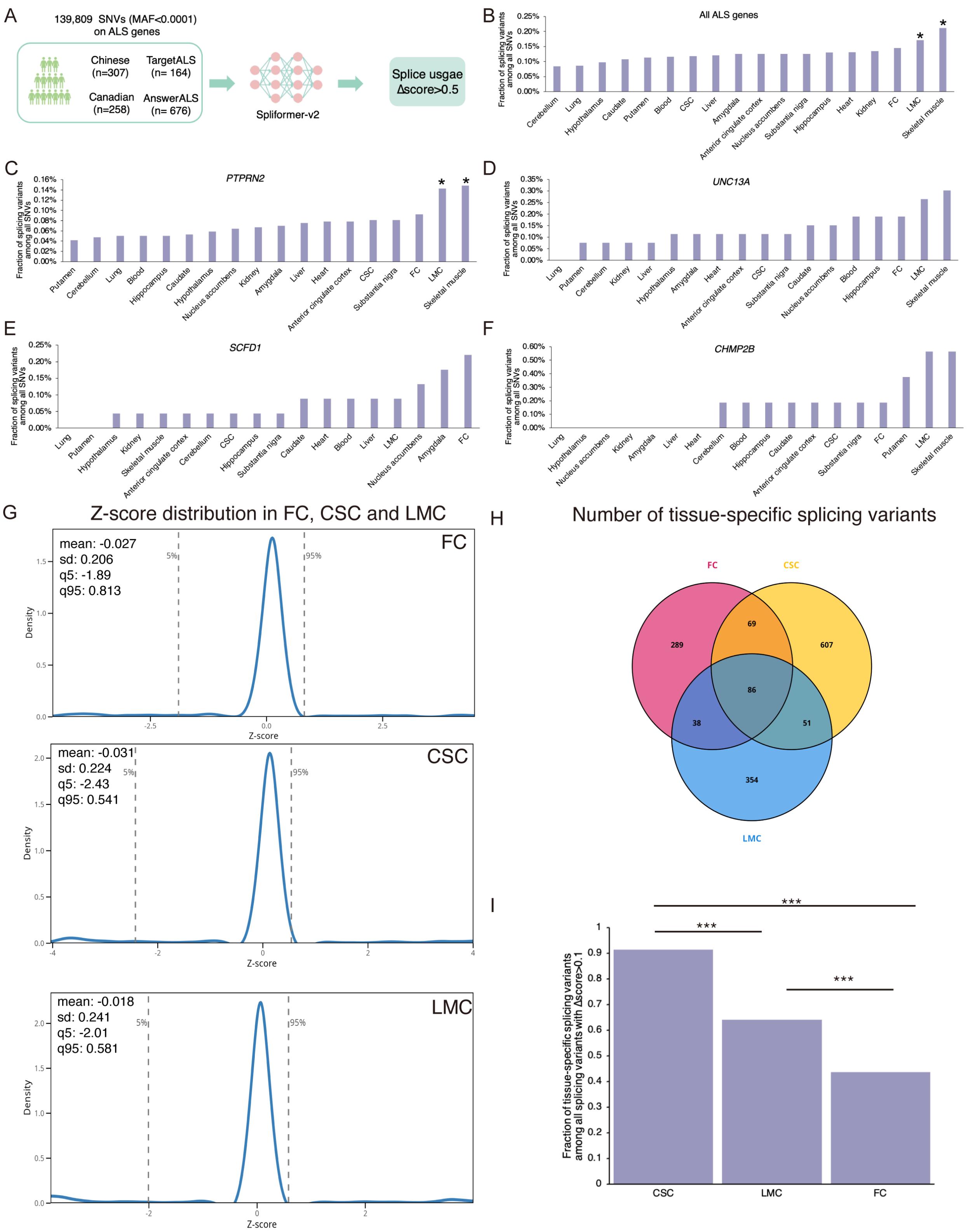


**Fig. S10 Tissue specific vulnerability to splicing variants in ALS. (A)** The workflow for identifying rare splicing variants in 18 tissues. 139,809 SNVs (AF<0.0001) were extracted from four ALS cohorts: Chinese (n=307), Canadian (n=258), TargetALS (n=164) and AnswerALS (n=676). SNVs with a predicted |Δscore| > 0.5 were classified as splicing variants. **(B)** Fraction of splicing variants across 18 tissues in four cohorts. **(C-F)** Fraction of splicing variants located on *PTPRN2* (**C**), *UNC13A* (**D**), *SCFD1*(**E**) and *CHMP2B* (**F**) across 18 tissues in four cohorts. **(G)** Distribution of Z-score in FC, CSC and LMC. A reference mutation set was constructed using 855 SNVs that exhibited similar splicing effects (|Δscore| > 0.1 in carriers vs. noncarriers for each tissue) and consistent directional changes in splicing usage across all three tissues. The Spliformer_V2 models (CSC, FC, and LMC) were used to predict the splicing effects of these variants, and the Δscore were used to calculate a Z-score distribution. This Z-score distribution serves as a reference for evaluating tissue specificity in subsequent analyses. **(H)** Venn diagram shows the number of tissue-specific splicing variants in FC, CSC and LMC. Tissue-specific splicing variants were defined as SNVs whose tissue z-score fell above than 95th percentile or below than 5th percentile of the reference distribution. **(I)** Bar figure shows the fraction of tissue-specific splicing variants among all splicing variants with |Δscore|>0.1 in CSC, FC and LMC. Fisher's exact test was used to assess the enrichment of tissue-specific splicing variants. $^{*}$ denotes as P_FDR<0.05 in (**B**). $^{***}$ denotes as P<0.0001 in (**I**).

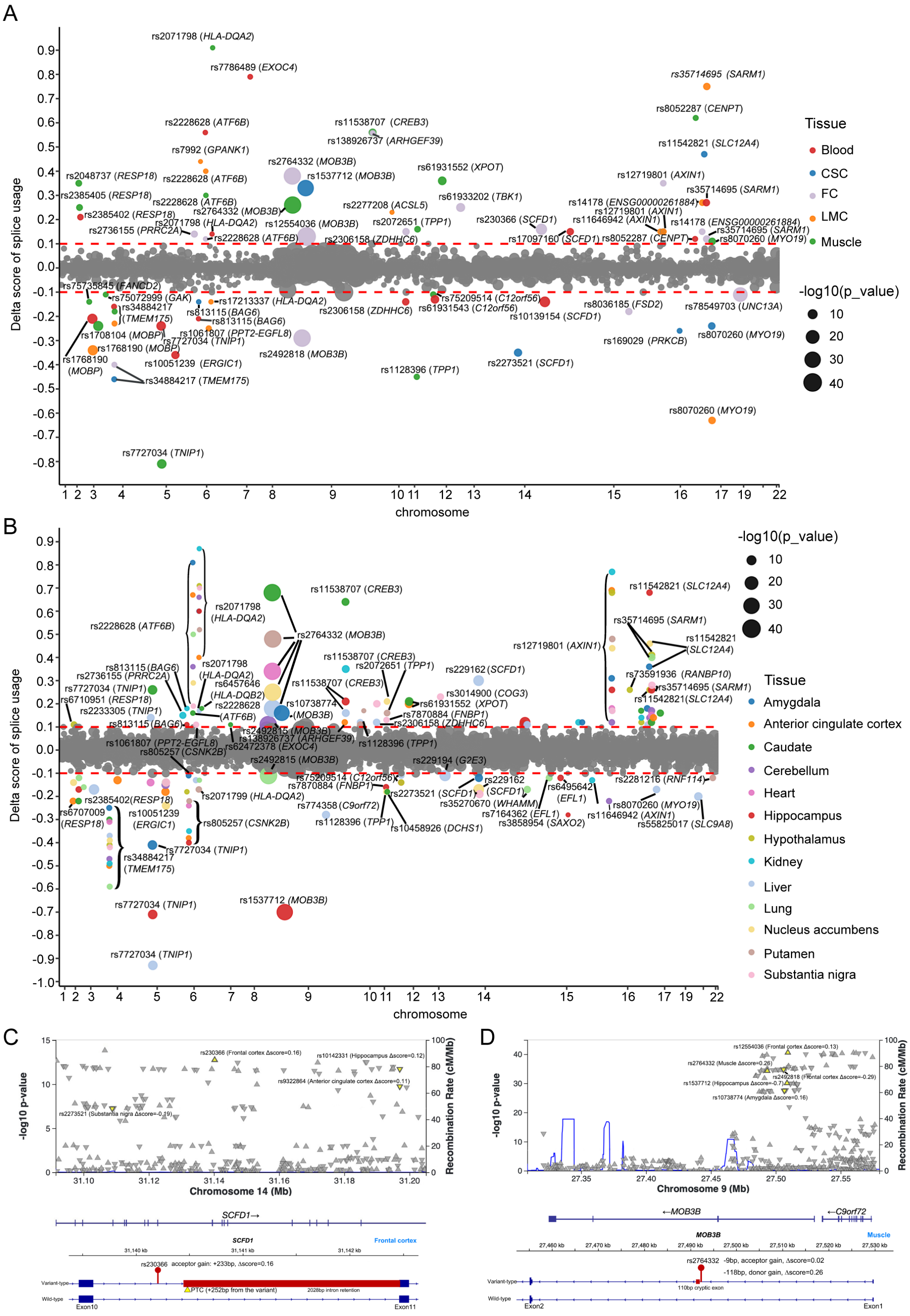
A
Delta score of splice usage
chromosome
Tissue
Blood
CSC
FC
LMC
Muscle
-log10(p_value)
10
20
30
40
rs2071798 (HLA-DQA2)
rs7786489 (EXOC4)
rs35714695 (SARM1)
rs8052287 (CENPT)
rs11538707 (CREB3)
rs138926737 (ARHGEF39)
rs2228628 (ATF6B)
rs7992 (GPANK1)
rs11542821 (SLC12A4)
rs2764332 (MOB3B)
rs1537712 (MOB3B)
rs61931552 (XPOT)
rs2048737 (RESP18)
rs2385405 (RESP18)
rs2385402 (RESP18)
rs2736155 (PRRC2A)
rs12554036 (MOB3B)
rs2306158 (ZDHHC6)
rs2277208 (ACSL5)
rs2072651 (TPP1)
rs61933202 (TBK1)
rs230366 (SCFD1)
rs17097160 (SCFD1)
rs12719801 (AXIN1)
rs11646942 (AXIN1)
rs14178 (ENSG00000261884)
rs8070260 (MYO19)
rs75735845 (FANCD2)
rs75072999 (GAK)
rs34884217 (TMEM175)
rs813115 (BAG6)
rs17213337 (HLA-DQA2)
rs1061807 (PPT2-EGFL8)
rs7727034 (TNIP1)
rs1708104 (MOBP)
rs1768190 (MOBP)
rs10051239 (ERGIC1)
rs2492818 (MOB3B)
rs75209514 (C12orf56)
rs61931543 (C12orf56)
rs10139154 (SCFD1)
rs8036185 (FSD2)
rs78549703 (UNC13A)
rs169029 (PRKCB)
rs2273521 (SCFD1)
rs1128396 (TPP1)
B
Tissue
Amygdala
Anterior cingulate cortex
Caudate
Cerebellum
Heart
Hippocampus
Hypothalamus
Kidney
Liver
Lung
Nucleus accumbens
Putamen
Substantia nigra
rs2071798 (HLA-DQA2)
rs2764332 (MOB3B)
rs11538707 (CREB3)
rs2072651 (TPP1)
rs229162 (SCFD1)
rs3014900 (COG3)
rs61931552 (XPOT)
rs7870884 (FNBP1)
rs2306158 (ZDHHC6)
rs6457646 (HLA-DQB2)
rs10738774 (MOB3B)
rs2492815 (MOB3B)
rs62472378 (EXOC4)
rs6710951 (RESP18)
rs2233305 (TNIP1)
rs805257 (CSNK2B)
rs229194 (G2E3)
rs12719801 (AXIN1)
rs73591936 (RANBP10)
rs2281216 (RNF114)
rs6495642 (EFL1)
rs7164362 (EFL1)
rs3858954 (SAXO2)
rs35270670 (WHAMM)
rs10458926 (DCHS1)
rs774358 (C9orf72)
rs2071799 (HLA-DQA2)
rs6707009 (RESP18)
rs1537712 (MOB3B)
rs55825017 (SLC9A8)
C
-log10 p-value
Recombination Rate (cM/Mb)
Chromosome 14 (Mb)
rs230366 (Frontal cortex Δscore=0.16)
rs10142331 (Hippocampus Δscore=0.12)
rs9322864 (Anterior cingulate cortex Δscore=0.11)
rs2273521 (Substantia nigra Δscore=-0.19)
SCFD1→
SCFD1
Frontal cortex
rs230366 acceptor gain: +233bp, Δscore=0.16
Variant-type
Wild-type
PTC (+252bp from the variant)
2028bp intron retention
Exon10
Exon11
D
-log10 p-value
Recombination Rate (cM/Mb)
Chromosome 9 (Mb)
rs12554036 (Frontal cortex Δscore=0.13)
rs2764332 (Muscle Δscore=0.26)
rs2492818 (Frontal cortex Δscore=-0.29)
rs1537712 (Hippocampus Δscore=-0.7)
rs10738774 (Amygdala Δscore=0.16)
←MOB3B
←C9orf72
MOB3B
Muscle
rs2764332 -9bp, acceptor gain, Δscore=0.02
-118bp, donor gain, Δscore=0.26
110bp cryptic exon
Variant-type
Wild-type
Exon2
Exon1

**Fig. S11 Predicted splicing effects of disease-associated GWAS hits ($p < 10^{-5}$) in ALS. (A, B)** Overview of predicted splicing effects in 18 tissues. The x-axis represents chromosomes, and the y-axis represents Δscore of splice usage predicted by Spliformer-V2 across different tissues. Colors denote different tissues, and point size corresponds to $-\log_{10}$(p-value). SNVs with labels indicate variants with |Δscore| > 0.1. （**C, D**) Fine-mapping plots highlighting ALS-associated GWAS variants in *SCFD1* (**C**) and *MOB3B* (**D**) and their predicted effects on RNA splicing. The *SCFD1* variant, rs230366, is predicted to increase the usage of an acceptor site, resulting in a 2,028-bp extension of exon 11 and the introduction of a PTC. The *MOB3B* variant, rs2764332, is predicted to increase the usage of both a donor and an acceptor site, leading to the inclusion of a 110-bp cryptic exon. SNVs are indicated by pins, exons by boxes, and PTCs by triangles. The variant location coordinates are based on hg19.

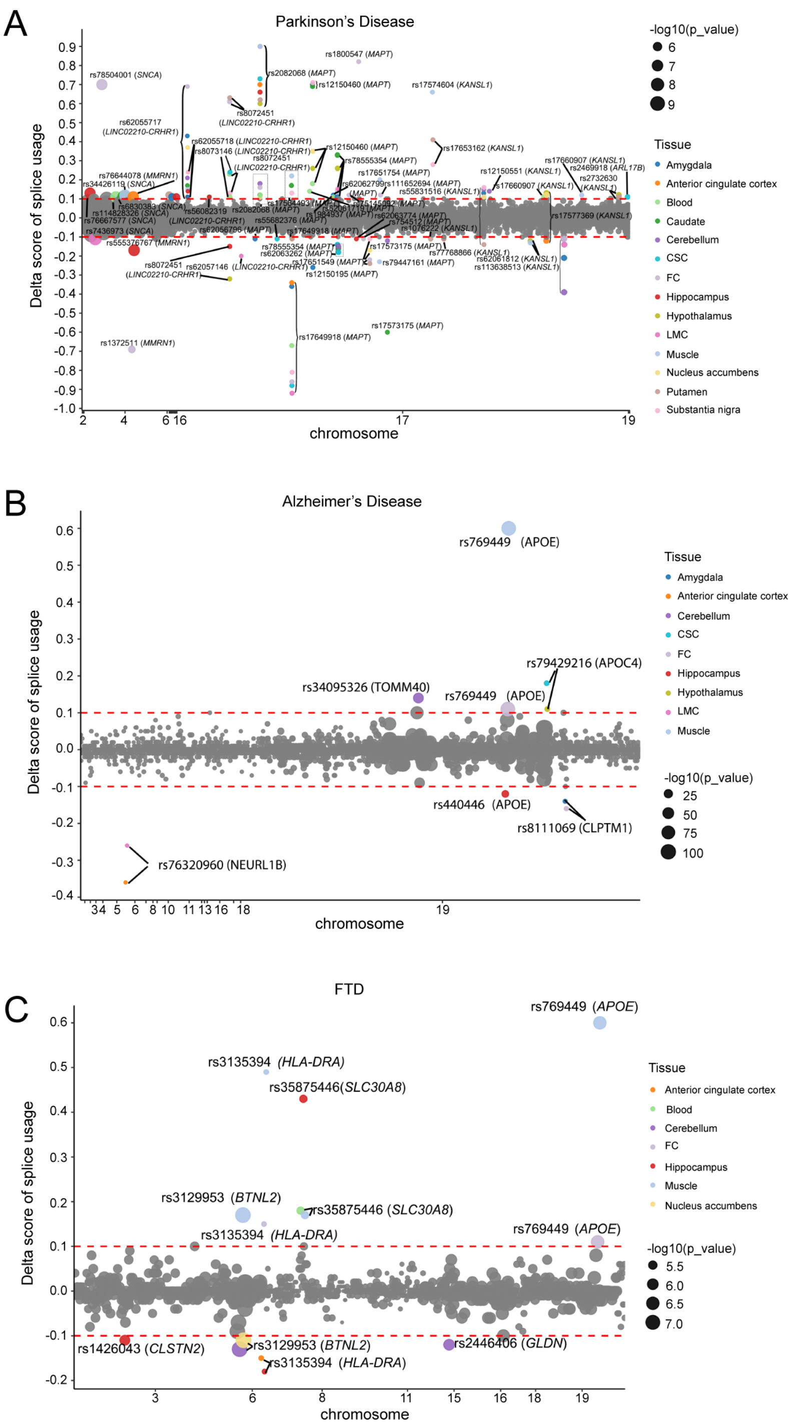


**Fig. S12 Predicted splicing effects of PD, AD and FTD-associated GWAS hits ($p < 10^{-5}$).** Dot-plots showing predicted splicing effects of PD **(A)**, AD **(B)** and **(C)** FTD associated variants in different tissues. The x-axis represents chromosomes, and the y-axis represents Δscore predicted by Spliformer-V2 across different tissues. Colors denote different tissues, and point size corresponds to $-\log_{10}$(p-value). SNVs with labels indicate variants with |Δscore| > 0.1. The variant location coordinates are based on hg19.

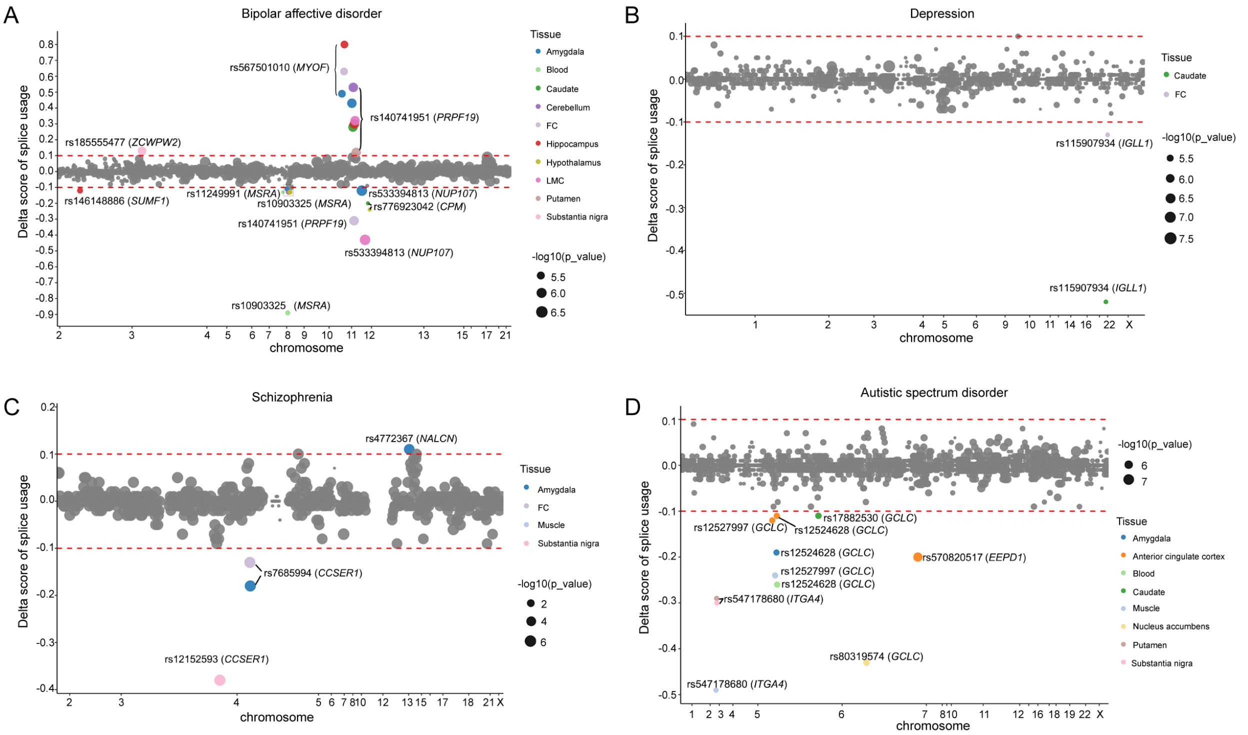


**Fig. S13 Predicted splicing effects of Bipolar affective disorder, Depression, Schizophrenia and ASD-associated GWAS hits ($p < 10^{-5}$).** Dot-plots showing predicted splicing effects of Bipolar affective disorder **(A)**, Depression **(B),** Schizophrenia **(C),** and Autistic spectrum disorder **(D)** associated variants in different tissues. The x-axis represents chromosomes, and the y-axis represents ΔSplice site usage values predicted by Spliformer-V2 across different tissues. Colors denote different tissues, and point size corresponds to $-\log_{10}$(p-value). SNVs with labels indicate variants with |Δscore| > 0.1. The variant location coordinates are based on hg19.

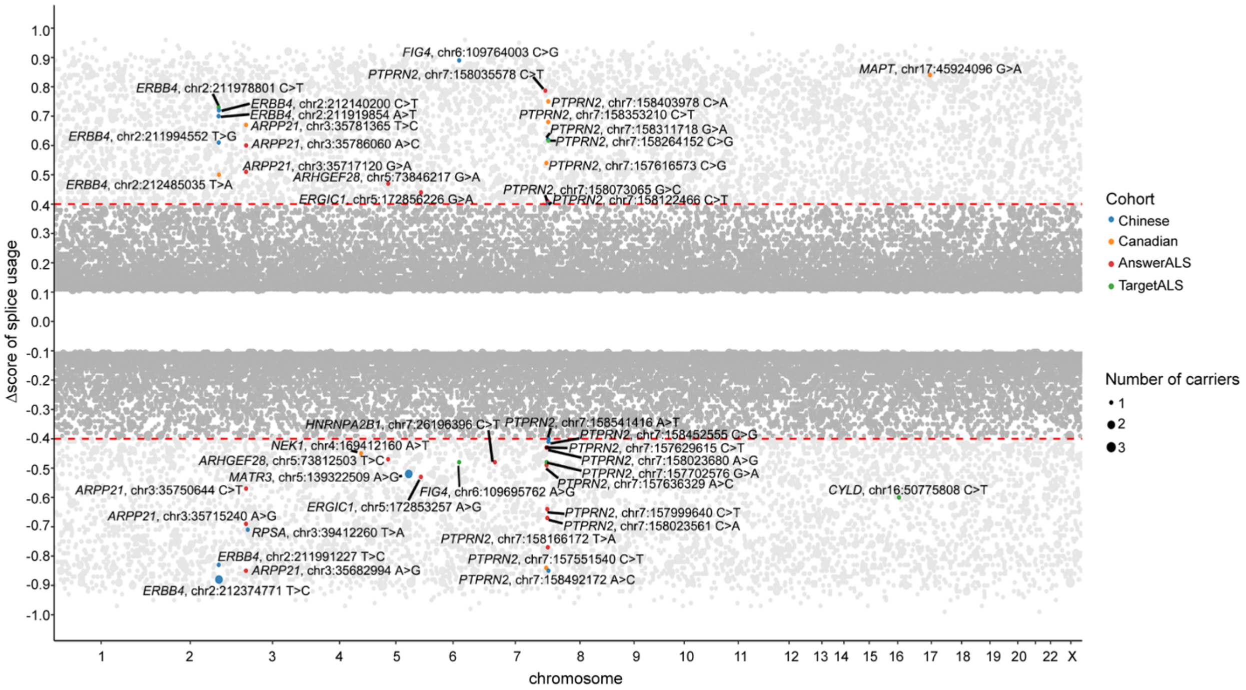


**Fig. S14 Predicted splicing effects of rare intronic variants in 1,405 ALS from Chinese (n=307), Canadian (n=258), TargetALS (n=164) and AnswerALS (n=686) datasets.** The x-axis represents chromosomes, and the y-axis represents Δscore predicted by Spliformer-V2 in LMC. Point size corresponds to carrier numbers. ALS gene names were labeled in the figure. The variant location coordinates are based on hg38.

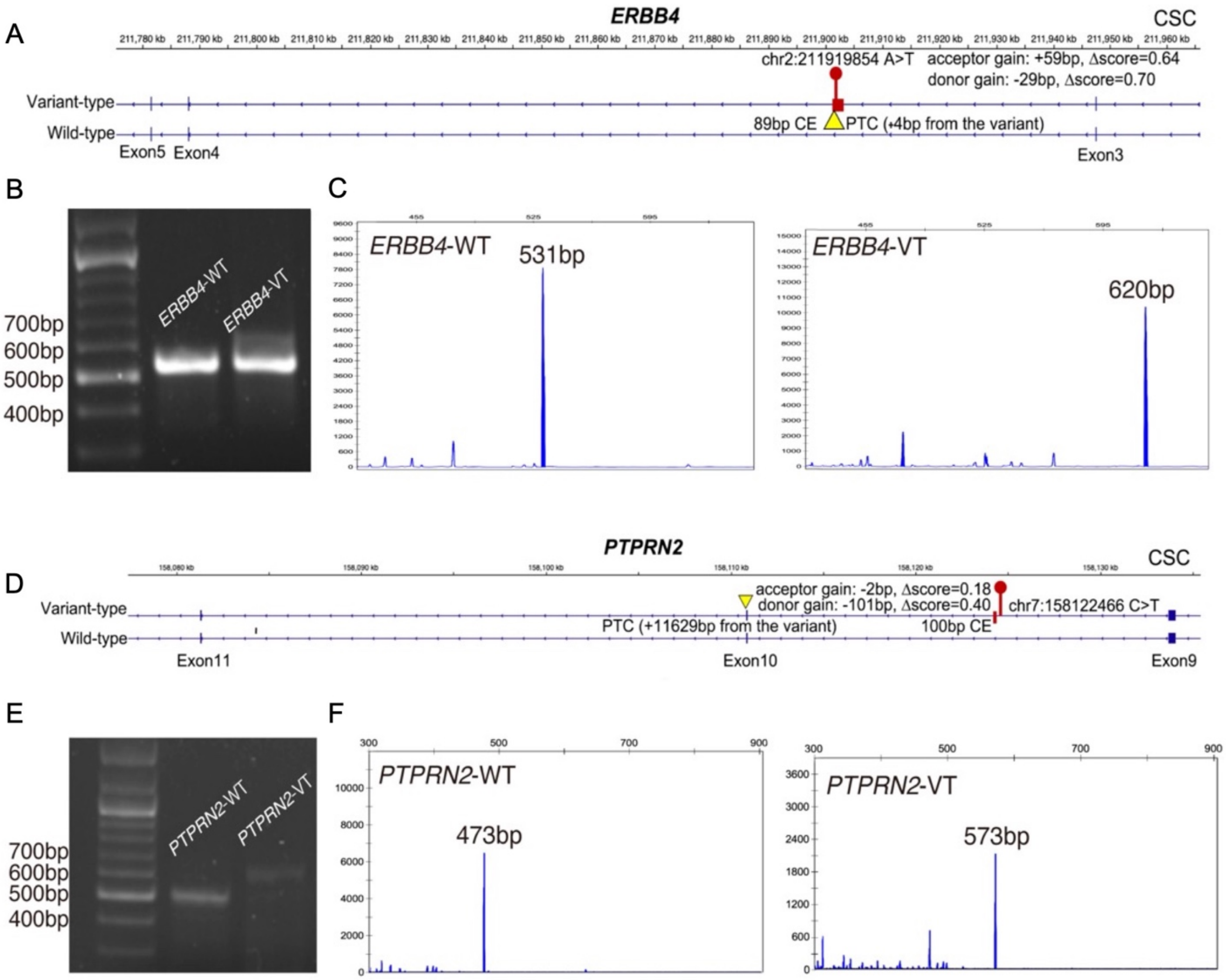


**Fig S15 (A)** Demonstration of RNA-splicing linked to the *ERBB4* variant (chr2:211919854 A>T, hg38), which is expected to cause an 89bp CE and introduce a PTC 4bp downstream of the variant. (**B**) Imaging of agarose gel electrophoresis of *ERBB4* RT-PCR products. (**C**) Amplicon length analysis of *ERBB4* RT-PCR products. The wild-type and variant-expressed sh-sy5y cells had different *ERBB4* mRNA transcripts (531 versus 620 bp). **(D)** Demonstration of RNA-splicing linked to the *PTPRN2* variant (chr7:158122466 C>T, hg38), which is expected to cause a 100bp CE and introduce a PTC 11,629 bp downstream of the variant. (**E)** Imaging of agarose gel electrophoresis of *PTPRN2* RT-PCR products. (**F**) Amplicon length analysis of *PTPRN2* RT-PCR products. The wild-type and variant-expressed sh-sy5y cells had different *PTPRN2* mRNA transcripts (473 versus 573 bp).

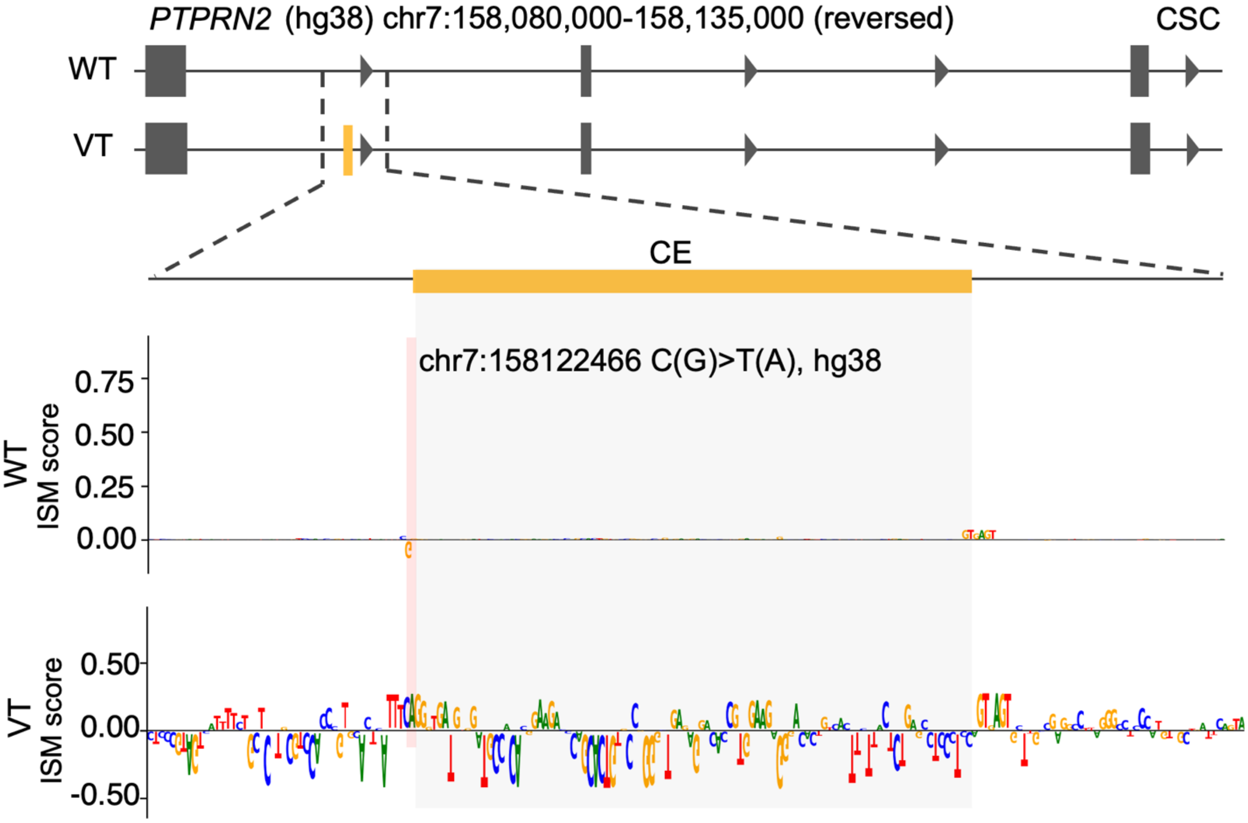


**Fig. S16 In silico mutagenesis analysis reveals splicing regulatory signals of the *PTPRN2* variant.** ISM profiles at the *PTPRN2* locus showing the variant induced cryptic exon (CE) inclusion. The variant rs555566959 (chr7:158122466 C>T, hg38) introduces a novel splice acceptor site, leading to CE inclusion in CSC. The top panel shows the local gene structure (hg38 coordinates), with the analyzed region highlighted and shown in a zoomed-in view below. Yellow boxes indicate the predicted cryptic exon or retained intronic region. The lower panels display in silico mutagenesis (ISM) profiles for the wild-type (WT) and variant (VT) sequences. Vertical pink lines indicate the position of the variant.

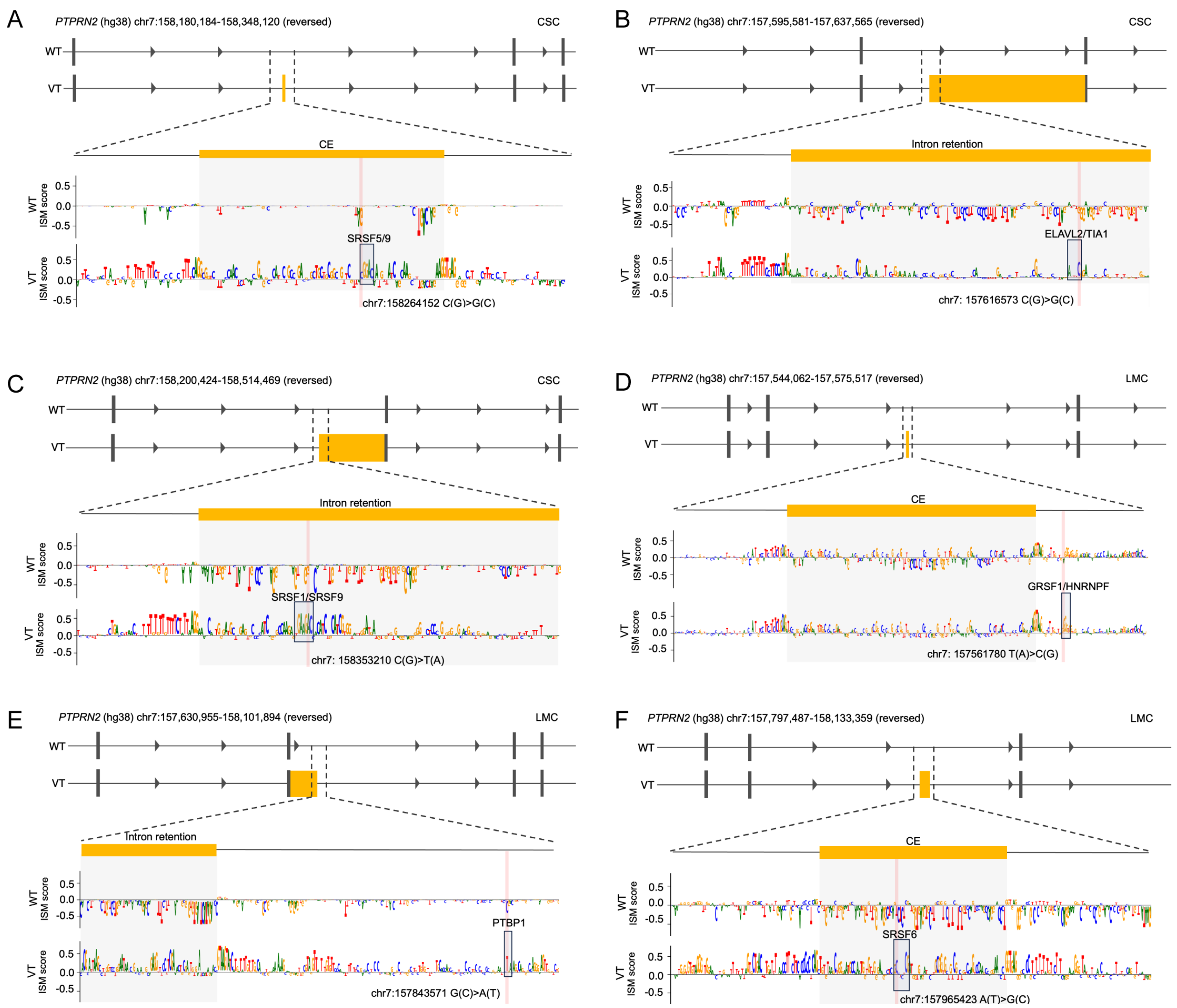


**Fig. S17 In silico mutagenesis analysis reveals splicing regulatory motifs at *PTPRN2* locus. (A–F)** Representative *PTPRN2* variants predicted to induce cryptic exon (CE) inclusion or intron retention (IR). The top panel shows the local gene structure (hg38 coordinates), with the analyzed region highlighted and shown in a zoomed-in view below. Yellow boxes indicate the predicted cryptic exon or retained intronic region. The lower panels display in silico mutagenesis (ISM) profiles for the wild-type (WT) and variant (VT) sequences. RBP-binding motifs potentially created by the variants are highlighted, including SRSF5/9 **(A)**, ELAVL2/TIA1 **(B)**, SRSF1/SRSF9 **(C)**, GRSF1/HNRNPF **(D)**, PTBP1 (**E**), and SRSF6 (**F**). Vertical pink lines indicate the positions of the variants.

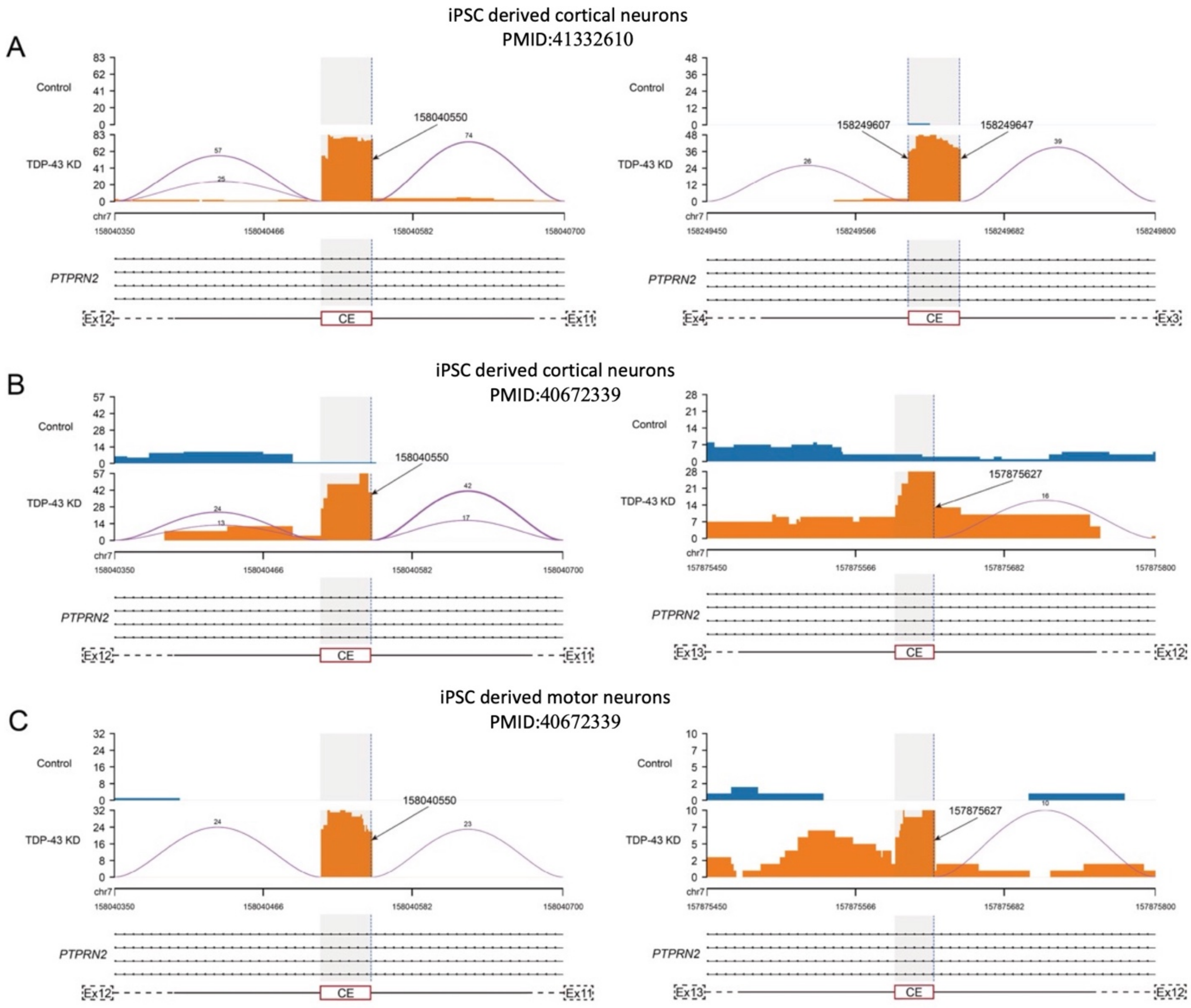


**Fig. S18 Sashimi plots showing representative *PTPRN2* cryptic splicing events upon TDP-43 knockdown in iPSC-derived cortical neurons (A-B); and iPSC-derived motor neurons (C).** The RNA-seq data were derived from previous publications (PMID:41332610; PMID:40672339). Coverage tracks represent RNA-seq read signals across the genomic regions, whereas arcs indicate detected splice junctions. Numbers above the arcs denote the numbers of junction-spanning reads supporting the corresponding splice junctions. Arrows and shaded regions indicate the candidate cryptic splice sites location (hg38 coordinates) and the surrounding regions of interest, respectively. Related to **Fig.5C, D, F**.

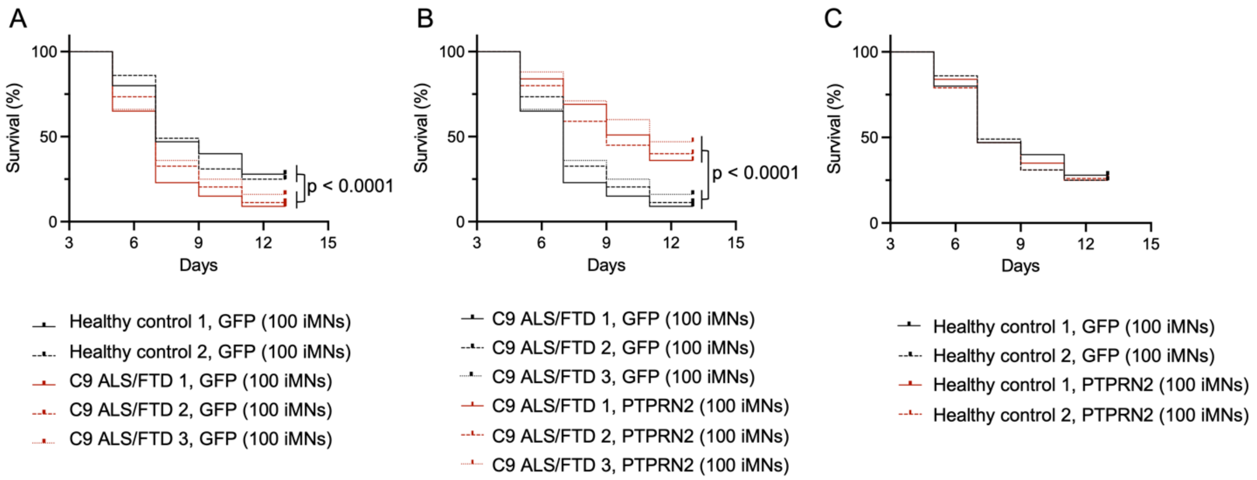


**Fig. S19 Kaplan–Meier curves showing survival status of iMNs from healthy controls or C9ORF72 patients.** Related to Figure 6G-I.

### Supplementary Tables

#### Supplementary Table 1 ISM-derived motifs and RBP associations in CSC-associated cryptic exon events

| Gene | Region | ISM score | RBP(s) | Motif feature | CNS-related |
|---|---|---|---|---|---|
| *ABLIM2* | acceptor | 0.34 | CELF4/5 | UG-rich | Yes, [1] |
| *AKAP11* | acceptor | 0.3 | PTBP1 | U-rich | no |
| *ANK1* | acceptor | 0.11 | ELAVL2 | U-rich | Yes, [2] |
| *ANXA7* | acceptor | 0.11 | CPEB2/4, U2AF2 | U-rich | no |
| *ARHGEF12* | acceptor | 0.32 | PTBP1 | U-rich | no |
| *ARHGEF12* | donor | 0.25 | PABPN1 | A-rich | no |
| *DDR1* | acceptor | 0.46 | PCBP2 | C-rich | no |
| *KCNT1* | acceptor | 0.25 | CELF4/5 | UG-rich | Yes, [1] |
| *KIF21A* | acceptor | 0.1 | ELAVL2 | U-rich | Yes, [2] |
| *LSM14B* | acceptor | 0.28 | ELAVL2 | U-rich | Yes, [2] |
| *MAP3K4* | acceptor | 0.28 | PTBP1 | U-rich | no |
| *MSRB3* | acceptor | 0.28 | ELAVL2 | U-rich | Yes, [2] |
| *MSRB3* | acceptor | 0.45 | U2AF2 | U-rich | no |
| *OTOF* | donor | 0.12 | PABPC5 | A-rich | no |
| *PHLDB1* | acceptor | 0.22 | ROD1 | U-rich | no |
| *PHLDB1* | donor | 0.14 | PCBP2 | C-rich | no |
| *PPP1R12A* | acceptor | 0.26 | TIA1 | U-rich | no |
| *RBFOX3* | acceptor | 0.26 | PCBP2 | C-rich | no |
| *SORBS1* | acceptor | 0.33 | hnRNPK | C-rich | no |
| *SYNE1* | acceptor | 0.45 | ELAVL2 | U-rich | Yes, [2] |
| *SYNE1* | donor | 0.6 | ZCRB1 | unclear | no |

| | | | | | |
|---|---|---|---|---|---|
| *TERF1* | acceptor | 0.21 | HNRNPCL1, U2AF2, HNRNPC | U-rich | no |
| *USP12* | acceptor | 0.32 | ELAVL2 | U-rich | Yes, [2] |

**Supplementary Table 2 Number of the variants in Clinvar dataset included in the Spliformer-v2 analyses**

| | Benign | Pathogenic |
|---|---|---|
| Intronic variants | 74,846 | 1,051 |
| Synonymous | 30,325 | 205 |
| All types | 105,171 | 1,256 |

**Supplementary Table 3 Sample characteristics of ALS patients and controls with whole genome sequencing data**

| | Chinese | Canadian | TargetALS | AnswerALS |
|---|---|---|---|---|
| **ALS** | | | | |
| Number of cases | 307 | 258 | 164 | 676 |
| Sex (n males, %) | 194 (63.2%) | 156 (60.5%) | 103 (62.8%) | 436 (64.5%) |
| Age at onset (y) | | | | |
| Mean (range) | 51 (19-84) | 58 (15-87) | 60 (28-81) | 57 (19-88) |
| Median (IQR) | 52 (42-62) | 59 (51-68) | 61 (54-68) | 58 (50-65) |
| Site of onset | | | | |
| Limb onset (n) | 266 | 126 | 104 | 464 |
| Bulbar onset (n) | 30 | 40 | 39 | 148 |
| Unknown | 11 | 92 | 21 | 64 |

IQR: interquartile range; y: years

**Supplementary Table 4. Information of rare splicing variants of *PTPRN2* in CSC and LMC**

| Gene | Variant (hg38) | AF | Prediction* | Result | Affected tissue |
|---|---|---|---|---|---|
| *PTPRN2* | chr7:157773597 G>A | . | acceptor gain: -12 bp, 0.72 | create MBNL1 motif | LMC |
| *PTPRN2* | chr7:157844167 C>G | 5.92E-05 | acceptor gain: +50 bp, 0.65 | create SRSF7 motif | LMC |
| *PTPRN2* | chr7:158122466 C>T | 7.23E-05 | acceptor gain: -2 bp, 0.92;<br>donor gain: -101 bp, 0.80 | directly create acceptor | LMC |

| | | | | | |
|---|---|---|---|---|---|
| *PTPRN2* | chr7:158403978 C>A | 1.31E-05 | acceptor gain: +116 bp, 0.81; donor gain: +6 bp, 0.55 | directly create donor | LMC |
| *PTPRN2* | chr7:157561780 T>C | . | acceptor gain: +194 bp, 0.51 | create GRSF1/HNRNPF motif | LMC |
| *PTPRN2* | chr7:158193154 C>T | 9.86E-05 | acceptor gain: -23 bp, 0.56 | create RBMX/SRSF9 motif | LMC |
| *PTPRN2* | chr7:158264152 C>G | 3.29E-05 | acceptor gain: +64 bp, 0.92; donor gain: -32 bp, 0.85 | create SRSF5/9 motif | LMC |
| *PTPRN2* | chr7:158190884 A>G | . | acceptor gain: -24 bp, 0.60 | create MBNL1 motif | LMC |
| *PTPRN2* | chr7:157592078 C>T | 3.29E-05 | acceptor gain: -184 bp, 0.63 | create SRSF1 motif | LMC |
| *PTPRN2* | chr7:157693642 C>G | 6.58E-06 | donor gain: -13 bp, 0.61 | create SRSF1 motif | LMC |
| *PTPRN2* | chr7:157843571 G>A | 5.92E-05 | donor gain: +169 bp, 0.67 | create PTBP1 motif | LMC |
| *PTPRN2* | chr7:157924091 C>G | 1.32E-05 | acceptor gain: +186 bp, 0.48 | create PCBP2/1 motif | LMC |
| *PTPRN2* | chr7:157965423 A>G | 6.57E-06 | acceptor gain: +42 bp, 0.62; donor gain: -59 bp, 0.50 | create SRSF6 motif | LMC |
| *PTPRN2* | chr7:158485253 C>G | 7.89E-05 | acceptor gain: -207 bp, 0.51 | create SRSF5 motif | LMC |
| *PTPRN2* | chr7:157616573 C>G | . | acceptor gain: +121 bp, 0.54 | create ELAVL2/TIA1 motif | CSC |
| *PTPRN2* | chr7:158122466 C>T | 7.23E-05 | acceptor gain: -2 bp, 0.18; donor gain: -101 bp, 0.4 | directly create acceptor | CSC |
| *PTPRN2* | chr7:158035578 C>T | 6.57E-05 | acceptor gain: +69 bp, 0.79; donor gain: -3 bp, 0.79 | directly create donor | CSC |
| *PTPRN2* | chr7:158073065 G>C | . | donor gain: +1 bp, 0.41 | directly create donor | CSC |
| *PTPRN2* | chr7:158264152 C>G | 3.29E-05 | acceptor gain: +64 bp, 0.53; donor gain: -32 bp, 0.62 | create SRSF5/9 motif | CSC |
| *PTPRN2* | chr7:158353210 C>T | . | acceptor gain: +44 bp, 0.68 | create SRSF1/9 motif | CSC |
| *PTPRN2* | chr7:158528233 A>C | 3.00E-04 | donor gain: +0 bp, 0.79 | directly create donor | CSC |

*Spliformer-V2 prediction results: affected splice site: distance from the variant, Δscore

**Supplementary Table 5. *PTPRN2* expression as cell type markers in human motor cortex based on snRNA-seq of 16 controls (syn51105515).**

| gene | p_val* | avg_log2FC | pct.1 | pct.2 | p_val_adj* | cluster |
|---|---|---|---|---|---|---|
| *PTPRN2* | 0 | 0.90051529 | 0.995 | 0.565 | 0 | Ex_L2_L3 |
| *PTPRN2* | 1.26E-257 | 1.3591921 | 0.996 | 0.593 | 4.61E-253 | Ex_L5 |
| *PTPRN2* | 2.23E-217 | 0.5445296 | 0.989 | 0.572 | 8.13E-213 | Ex_L4_L6 |
| *PTPRN2* | 0 | 0.79590589 | 0.997 | 0.57 | 0 | Ex_L3_L5 |
| *PTPRN2* | 6.01E-45 | 0.31305234 | 0.981 | 0.59 | 2.19E-40 | Ex_L4_L5 |
| *PTPRN2* | 0 | 1.22130354 | 0.998 | 0.581 | 0 | Ex_L6 |
| *PTPRN2* | 0 | 0.79554459 | 0.997 | 0.571 | 0 | Ex_L5_L6 |
| *PTPRN2* | 1.01E-286 | 1.25724487 | 0.925 | 0.591 | 3.69E-282 | In_SOM |
| *PTPRN2* | 5.64E-66 | 0.58737435 | 0.97 | 0.593 | 2.06E-61 | In_Rosehip |
| *PTPRN2* | 7.84E-154 | 0.73912309 | 0.906 | 0.589 | 2.86E-149 | In_5HT3aR |
| *PTPRN2* | 0 | 1.12178784 | 0.979 | 0.582 | 0 | In_PV |

* P-values were obtained from Wilcoxon rank-sum tests performed to compare each cell type against all other cell populations, with FDR correction applied.

**Supplementary Table 6 WGS/RNA-seq Data sources of 18 human tissues from the TargetALS and GTEx dataset**

| Tissue | Sample | Dataset |
|---|---|---|
| Amygdala | 20 controls | GTEx |
| Anterior cingulate cortex | 20 controls | GTEx |
| Caudate | 20 controls | GTEx |
| Cerebellum | 20 controls | GTEx |
| Frontal Cortex | 20 controls | GTEx |
| Hippocampus | 20 controls | GTEx |
| Hypothalamus | 20 controls | GTEx |
| Nucleus accumbens | 20 controls | GTEx |
| Putamen | 20 controls | GTEx |
| Substantia nigra | 20 controls | GTEx |
| Skeletal muscle | 20 controls | GTEx |
| Heart | 20 controls | GTEx |

| | | |
|---|---|---|
| Blood | 20 controls | GTEx |
| Kidney | 20 controls | GTEx |
| Liver | 20 controls | GTEx |
| Lung | 20 controls | GTEx |
| LMC | 10 controls and 10 ALS patients | TargetALS |
| CSC | 10 controls and 10 ALS patients | TargetALS |

**Supplementary Table 7 Primers used for RT-PCR experiments and amplicon length analyses in minigene assay**

| Gene | Forward Primer (5'-3') | Reverse Primer (5'-3') |
|---|---|---|
| *PTPRN2* | 6FAM-CCACTGTGCTGGATATCTGC | ACTGCTGGGCACCTGCAGGA |
| *ERBB4* | 6FAM-CGCAAATGGGCGGTAGGCGTG | CGTTCAATTGCCGACCCCTC |

**Supplementary Table 8 Information of siRNA or shRNA**

| Gene | Company | Information/Sequence |
|---|---|---|
| TARDBP siRNA | ThermoFisher Scientific | Cat #4427037, ID s530937 |
| PTPRN2 siRNA | ThermoFisher Scientific | Cat #4427037, ID s11567 |
| PTPRN2 shRNA | Tsingke Biotechnology | GATCGGACCAGCAGTGACCTTCAACTCG AGTTGAAGGTCACTGCTGGTCTTTTTT |

**Supplementary Table 9 Primers used for RT-qPCR experiments in SH-SY5Y cells**

| Gene | Forward Primer (5'-3') | Reverse Primer (5'-3') |
|---|---|---|
| *PTPRN2* | TTTACCGCTACGAGGTGTCG | TTCGGGAGGTCTGCAAGTTC |
| *GAPDH* | GACAGTCAGCCGCATCTTCT | GCGCCCAATACGACCAAATC |

**Supplementary Table 10 Primers used for iMNs experiments**

| Oligonucleotides and antibodies | Source | Sequence |
|---|---|---|
| qPCR for human *TARDBP*-43_F | Melamed et al. [3] | TCATCCCCAAGCCATTCAGG |
| qPCR for human *TARDBP*-43_R | Melamed et al. [3] | TGCTTAGGTTCGGCATTGGA |
| qPCR for human *PTPRN2*_F | Origene | TTCTCGGACCAGCAGTGACCTT |
| qPCR for human *PTPRN2*_R | Origene | TCAGTCCAGAGGTTTCCTCCAG |
| qPCR for human *GAPDH*_F | This paper | TGGTATCGTGGAAGGACTCATG |
| qPCR for human *GAPDH*_R | This paper | AGTAGAGGCAGGGATGATGTTC |

**Supplementary Table S11 Information of iPSC Cell lines**

| Cell line | Source | Identifier |
|---|---|---|
| Human: HC iPSC 1 | NINDS Biorepository | ND03231 |
| Human: HC iPSC 2 | NINDS Biorepository | ND05280 |
| Human: C9 ALS/FTD iPSC 1 | NINDS Biorepository | ND06769 |
| Human: C9 ALS/FTD iPSC 2 | NINDS Biorepository | ND12099 |
| Human: C9 ALS/FTD iPSC 3 | NINDS Biorepository | ND10689 |
| Human: dCas9-BFP-KRAB + TetO-Ngn2 iPSC | Kampmann lab [4] | WTC11 |

#### Supplementary Files

**Supplementary File 1:** Spliformer-V2 prediction results of GWAS indicated ALS associated variants (Δscore$>0.1$)

**Supplementary File 2:** Spliformer-V2 prediction results of GWAS indicated AD, PD, FTD associated variants (Δscore$>0.1$)

**Supplementary File 3:** Spliformer-V2 prediction results of GWAS indicated bipolar, schizophrenia, autism and depression associated variants (Δscore$>0.1$)

#### Supplementary Reference


1. Dasgupta, T. & Ladd, A.N. The importance of CELF control: molecular and biological roles of the CUG-BP, Elav-like family of RNA-binding proteins. *Wiley Interdiscip Rev RNA* **3**, 104-121 (2012).
2. Moakley, D.F. et al. Reverse engineering neuron-type-specific and type-orthogonal splicing-regulatory networks using diverse cellular transcriptomes. *Cell Rep* **44**, 115898 (2025).
3. Melamed, Z. et al. Premature polyadenylation-mediated loss of stathmin-2 is a hallmark of TDP-43-dependent neurodegeneration. *Nat Neurosci* **22**, 180-190 (2019).

4. Tian, R. et al. CRISPR Interference-Based Platform for Multimodal Genetic Screens in Human iPSC-Derived Neurons. *Neuron* **104**, 239-255 e212 (2019).
